# Persulfidation of DJ-1 : Mechanism and Consequences

**DOI:** 10.1101/2022.11.09.515774

**Authors:** Erwan Galardon, Nicolas Mathas, Dominique Padovani, Laurent Le Corre, Gabrielle Poncet, Julien Dairou

## Abstract

DJ-1 (also called PARK7) is a ubiquitously expressed protein involved in the etiology of Parkinson disease and cancers. At least one of its three cysteine residue is functionally essential, and its oxidation state determines the specific function of the enzyme. DJ-1 was recently reported to be persulfidated in mammalian cell lines, but the implications of this post-translational modification have not yet been analyzed. Here, we report that recombinant DJ-1 is reversibly persulfidated at cysteine 106 by reaction with various sulfane donors and subsequently inhibited. Strikingly, this reaction is orders of magnitude faster than C106 oxidation by H_2_O_2_, and persulfidated DJ-1 behaves differently than sulfinylated DJ-1. Both these PTM most likely play a dedicated role in DJ-1 signaling or protective pathways.

## 1. Introduction

DJ-1 (also called PARK7) is a small (~ 20 kDa) ubiquitously expressed homodimeric protein. Since the report that mutation of its encoding gene in humans leads to autosomal recessive early-onset Parkinson disease (PD) 7 [1], intensive studies have been undertaken to decipher its function and its role in the etiology of this neurodegenerative disease. Thus, DJ-1 has been proposed to take part in various physiological pathways related to the promotion of cell survival [2]. For instance, DJ-1 activates the extracellular signal-regulated kinase pathway ERK1/2 [3] and the phosphatidylinositol-3-kinase (PI3K/Akt) pathway [4]. It also modulates oxidative and electrophilic stresses. For example, DJ-1 activates the Nrf2-mediated antioxidant response [5], catalytically protects various biomolecules against glycation by methylglyoxal [6] or detoxifies reactive compounds produced during glycolysis [7,8], although the nature of its physiological substrates is still a matter of controversy [9,10]. In addition to PD, DJ-1 is now proposed to be involved in various pathological settings, like ischemia–reperfusion injury [11], inflammatory bowel disease [12], diabetes [13] or cancers [14].

Human DJ-1 is characterized by an α/β-flavodoxin fold core and it possesses three cysteines (C46, C53 and C106), the latter being highly conserved and localized in the nucleophile elbow region. C106 has been identified as an important residue whose thiolate group is key to most of DJ-1 functions [5], its oxidation state determining the specific function of the enzyme. Thus, under a reduced catalytic C106 status (CysS^-^) [5], DJ-1 has been proposed to act as a peroxiredoxin-like peroxidase [15], a protease [16], a glyoxalase [8], a deglycase [17] and more recently as a scavenger of a reactive glycolytic metabolite [7]. Additionally, DJ-1 displays a non-physiologically relevant esterase-activity which was used to develop an assay to screen for new DJ-1 inhibitors [18]. On another hand, C106 is characterized by a low thiol p*K_a_* value ~ 5 [19] and can be oxidized to sulfinate (-SO_2^-^_) and sulfonate (-SO_3^-^_) by hydrogen peroxide (H_2_O_2_) [20,21]. For instance, the sulfinylation of C106 shifts its isoelectric point and promotes its intracellular relocation, allowing DJ-1 to play a role in redox sensing and cytoprotection [22,23]. The same post-translational modification (PTM) regulates its participation in the composition of high molecular weight complexes that play a role in RNA metabolism and catecholamine homeostasis in cultured cells and human brain [24,25]. Interestingly, whilst C106 was clearly demonstrated to be the key cysteine in the aforementioned studies, the sulfinylation of C46 may also have physiological significance as the protein thus modified is one of the few substrates of sulfiredoxin [26].

In addition to being implicated into the various S-oxygenation reactions briefly described above, C106 is also the target of other PTMs. For instance, all three Cys residues of DJ-1 were reported to be nitrosylated (formation of CysS-NO) in various cell lines in conflicting studies [27,28], with C106 seemingly playing a role in trans-nitrosylation processes [28,29]. In addition, C106 was also found to experience persulfidation (formation of CysS-SH), which may prevent it from uncontrolled S-oxygenation under oxidative stress conditions in MEF cells [30]. In addition to the importance of C106 as a redox sensor, part of the activity of DJ-1 depends on the redox-sensitive removal of a 15-amino acid peptide at its C terminus (C_ter_) [16,31]. For instance, the C_ter_ cleavage of DJ-1 in response to acute myocardial ischemia-reperfusion injury protects from heart failure by inducing anti-glycation properties [32]. However, the nature of the stimulus and the mechanism by which cleavage of the C_ter_ peptide occurs are still elusive. In this context, understanding the reactivity and the effects of various redox messengers on DJ-1 might provide new insights into DJ-1 role in various cellular contexts and could identify novel mechanisms involved in diseases’ setting and etiology.

Here, we report our first results on the persulfidation of DJ-1, and its outcome on the enzymatic activities and structure of the protein. We show that recombinant human DJ-1 is persulfidated at C106 and inhibited *in vitro* by reaction with various sulfane sulfur [33] donors, a reaction orders of magnitude faster than sulfinylation. Additionally, recombinant DJ-1 is endogenously persulfidated when overexpressed in *E. coli*. Although persulfidation and sulfinylation both result in DJ-1 inhibition, they lead to proteins with different behavior. These observations suggest different fates for each of these PTMs.

## 2. Materials and Methods

### 2.1 Materials

Most chemical and biochemical reactants were purchased from Merck. Sodium di- and tretrasulfide were purchased from Dojindo Molecular Technologies Inc. and sodium hydrosulfide from Strem. These salts were manipulated under argon atmosphere (< 1 ppm O_2_) in a glovebox. DAz-2:Cy5 was synthesized as previously described[30]. Recombinant human thioredoxin 1 (hTrx) was purchased from ThermoFisher. The plasmid pET-TRSter for heterologous expression of human thioredoxin reductase (hTrxR) was purchased from Addgene. Plasmids for wt and mutant DJ-1 were obtained from Dr. Sun-Sin Cha[34]. The plasmid containing the human CSE gene (pET-28-based expression vector incorporating a tobacco etch virus (TEV)-cleavable N-terminal His tag fusion) was a kind gift from Dr. Tobias Karlberg (Structural Genomics Consortium, Karolinska Institute, Sweden). Reactions were typically run in phosphate buffered saline (PBS) containing 200 μM diethylenetriaminepentaacetic acid (DTPA), unless otherwise stated. The buffer was roughly degassed by bubbling argon for 30 minutes before experiments with the sodium hydrosulfide or polysulfide salts. UV-Visible spectra were recorded on a Cary 300, Jasco V-700 or Biotek PowerWave XS spectrometers. Differential Scanning Fluorimetry (DSF) experiments were carried out on a Bio-rad CFX96 Real Time PCR system. Gels were imaged on a LAS 4000 (GE Healthcare) or Bio-Rad GelDoc Go and images were treated with FiJi. The Liquid Chromatography coupled to Mass Spectrometry (LC-MS) system was composed of Shimadzu apparatus equipped with a LC30AD pump and a kinetex 5u C18 100A column, a SiL30AC auto-sampler coupled with a photodiode array detector PDA20A and a triple quadrupole mass detector 8060. Fitting of the data was performed with SigmaPlot 10 (Systat Software). Statistical analysis was carried out using the Excel (Microsoft) data analysis package: Activities (Figure 2 & 3) or persulfidation levels (Figure 5) were compared with the relevant controls using unpaired t-test.

**Figure 1.**
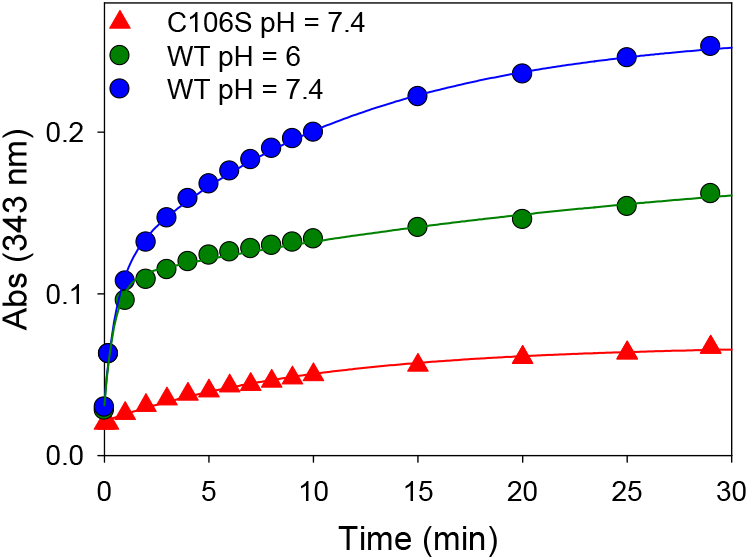
Reactivity of wt or mutant C106S DJ-1 with 2,2’-dithiopyridine. Time course of thiopyridine release monitored at 343 nm upon addition of DTP (180 μM final) to an 18 μM solution of wt DJ-1 or its mutant C106S at 25°C in PBS (pH = 7.4) or MES buffer (50 mM, pH=6.0) containing 0.2 mM DTPA.

**Figure 2.**
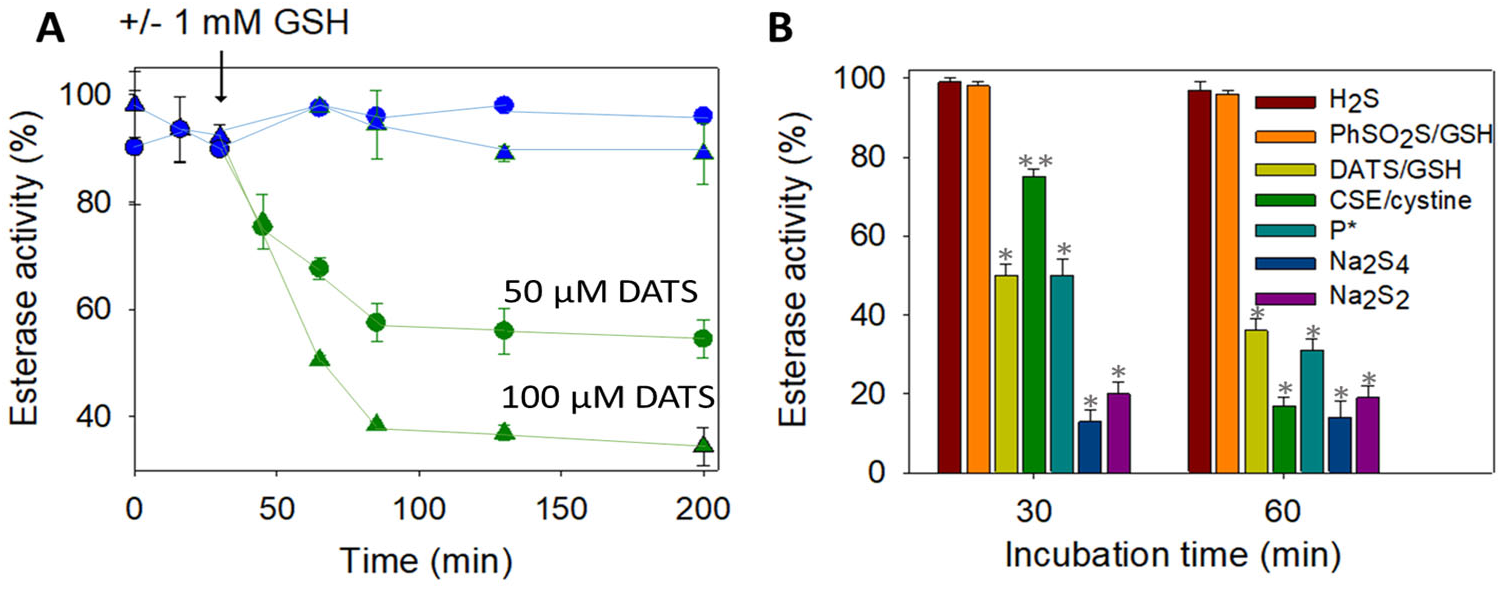
**(A) Wt DJ-1 activity is inhibited by DATS/DTT** – Time-dependence of DJ-1 esterase activity (reported as a % of the hydrolytic activity of purified DJ-1, determined as described in the experimental section). 12.5 μM DJ-1 was incubated at 25°C in PBS buffer with 50 μM (circles) or 100 μM (triangles) DATS with (green) or without (blue) 1 mM GSH added at t = 30 minutes. GSH alone did not show significant hydrolase activity (less than 10%). **(B) Wt DJ-1 activity is inhibited by sulfane donors** – Incubation of DJ-1 (12.5 μM) with various sulfur-containing molecules for 30 or 60 minutes in PBS (H_2_S: 50 μM, DATS/GSH: 100 μM / mM, GSH/PhSO_2_SNa; 2.5 mM/0.5 mM, CSE/cystine: 4 μM/1 mM, P*: 50 μM, Na_2_S_4_: 4 μM, Na_2_S_2_: 12.5 μM). Data are presented as means ± SE of three independent experiments. *p < 0.01 **p < 0.05 *vs* control (unpaired t-test).

**Figure 3.**
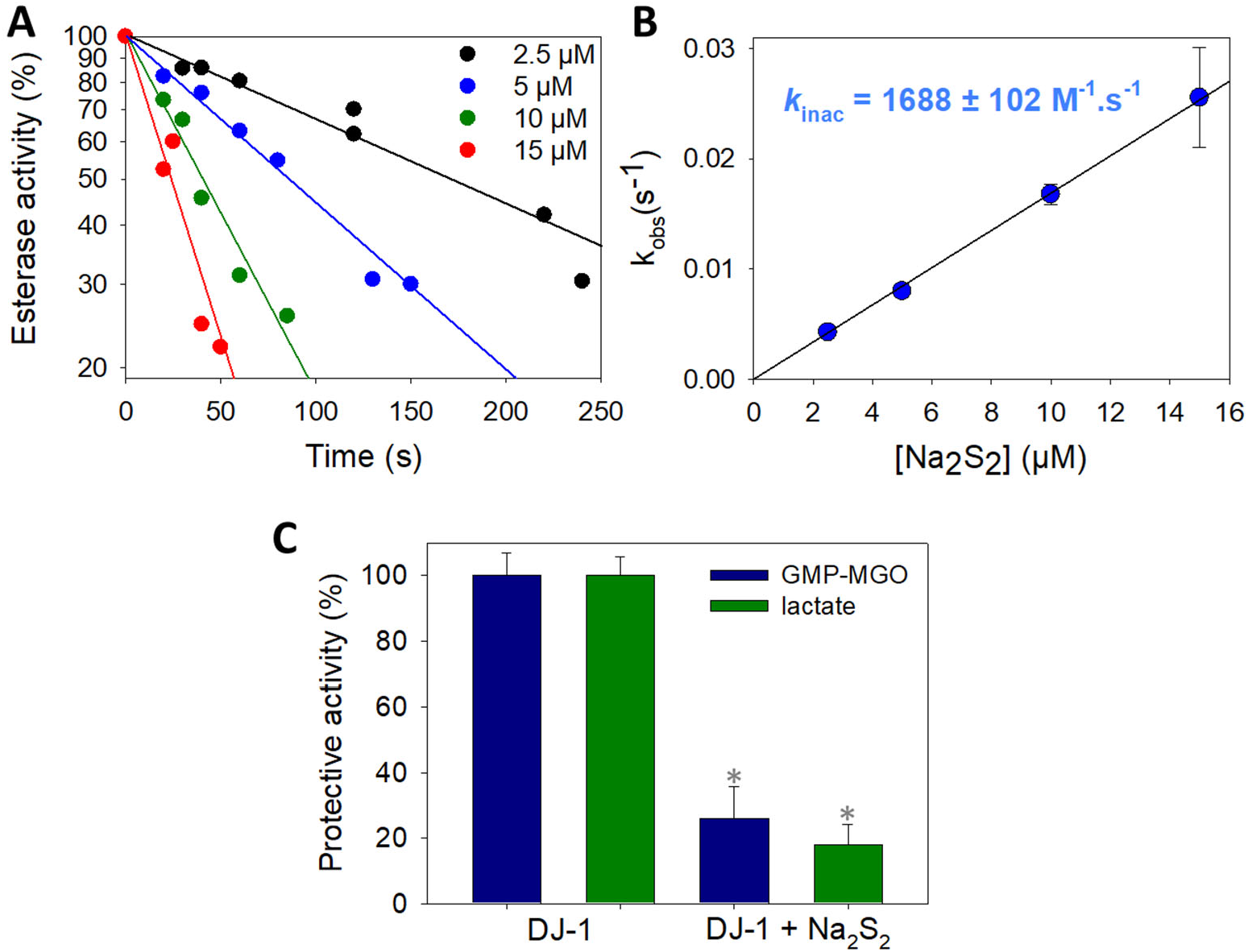
Kinetics of wt DJ-1 inactivation by sodium disulfide. (**A**) typical dataset obtained by recording DJ-1 (625 nM) activity at various time points after pre-incubation with various concentrations of Na_2_S_2_ (2.5, 5, 10 and 15 μM). (**B**) determination of the rate constant k_inac_ for the reaction between DJ-1 and Na_2_S_2_. Data are presented as means ± SE of three independent experiments. (**C**) **Protection against MGO-induced modification of GMP.** Protection against the formation of GMP-methylglyoxal adduct provided by wt DJ-1 or DJ-J1 treated with 1.5 equiv. Na_2_S_2_ calculated from the levels of glycated GMP (blue) or lactate (green) (for experimental details, see section 2.4). Data are presented as means ± SE of three independent experiments. *p < 0.01 (unpaired t-test).

**Figure 4.**
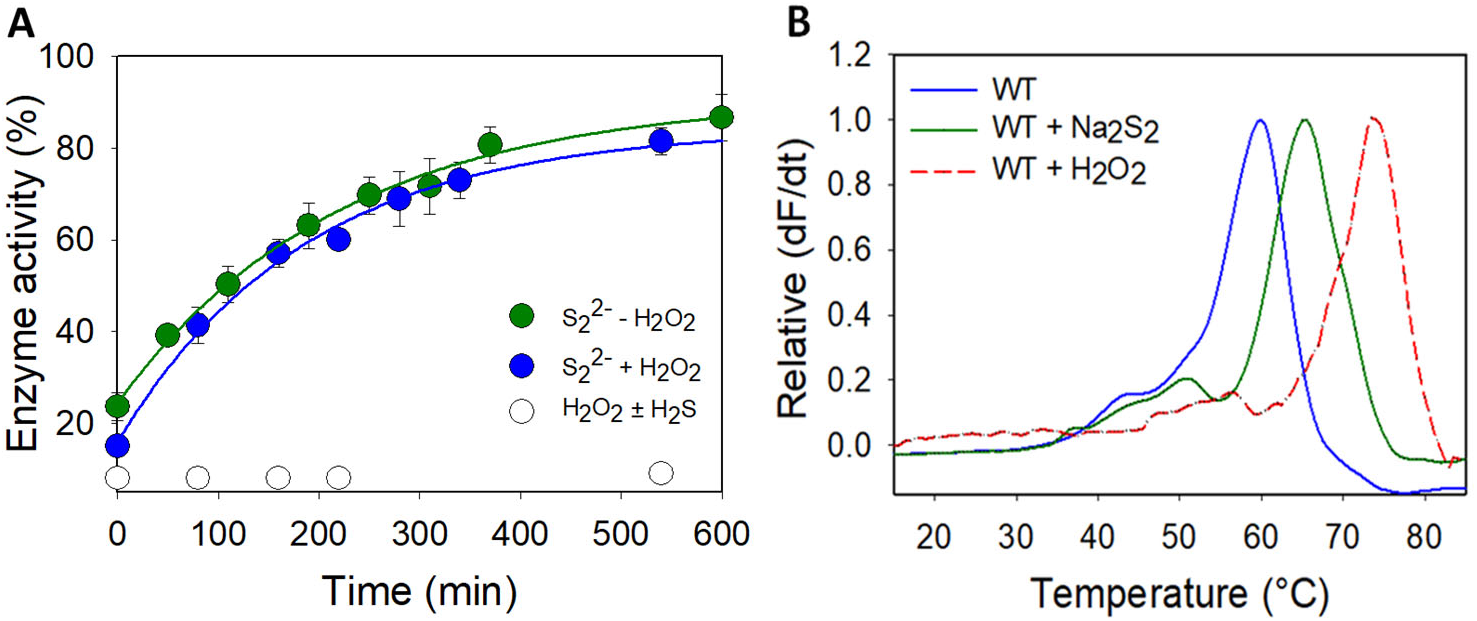
**(A) Reactivation of wt DJ-1 by DTT after sulfane sulfur treatment** - Esterase activity (reported as a % of the hydrolytic activity of purified DJ-1, determined as described in the experimental section) recorded over time after addition of DTT (1 mM) to: Green: a solution of inhibited DJ-1 obtained by incubating 12.5 μM DJ-1 with 12.5 μM Na_2_S_2_ for 20 min at RT. Blue: Inhibited DJ-1 was pre-incubated with hydrogen peroxide (80 μM, 30 min then 300 U of catalase, 15 min) before addition of 1 mM DTT. White: a 12.5 μM solution of inhibited DJ-1 obtained by incubating 50 μM DJ-1 with 150 μM H_2_O_2_ with/without 150 μM NaSH for 30 min at RT, and dilution. Data are presented as means ± SE of two independent experiments. **Figure (B) Wt DJ-1 is stabilized by sodium disulfide** - Differential scanning fluorimetry (ThermoFluor) spectra obtained for wt DJ-1 (blue), its inhibited form with Na_2_S_2_ (green), or its sulfinic form in the presence of DTT (red). Data are presented as representative of three independent experiments.

**Figure 5.**
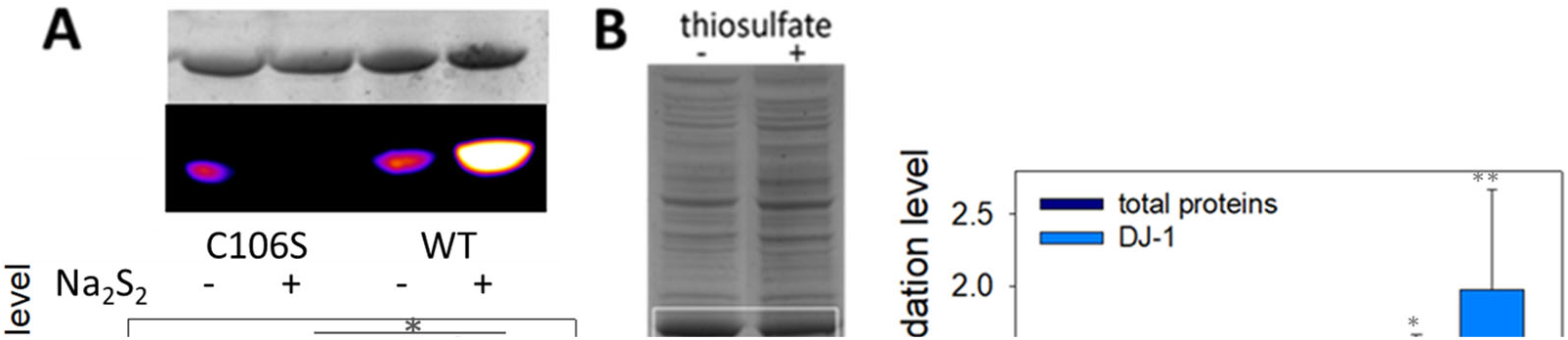
**(A) Wt DJ-1 is persulfidated at C106** – Representative in-gel fluorescence (Cy5-canal) and Coomassie Brilliant Blue staining scans obtained for the mutant C106S and wt DJ-1 with/without Na_2_S_2_, and relative persulfide level (Cy5/Coomassie) (n = 4, 70 μM DJ-1 ± 150 μM Na_2_S_2_, see the experimental section for full details) *p < 0.01 (unpaired t-test). **(B) Wt DJ-1 is persulfidated in *E. coli*** - Typical in-gel fluorescence scans (Cy5-canal and Coommassie Brilliant Blue staining) obtained after lysis of *E. coli* cultures supplemented with ± 25 mM thiosulfate. The signals corresponding to DJ-1 are circled in white. **(C) Effect of sulfane donors** - Relative persulfidation levels (calculated as the Cy5/CBB signal ratio) of total proteins (dark blue) and DJ-1 (light blue) in E coli lysates as a function of the sulfur-compound added to the culture medium (no additive, 1 mM cysteine trisulfide, 25 mM sodium thiosulfate). Data are presented as means ± SE of at least 3 independent experiments, *p < 0.01 **p < 0.*05 vs* experiments without donor (unpaired t-test).

### 2.2. Proteins Expression and Purification

Proteins were expressed and purified as previously described[17], with the exception that the last purification step of DJ-1, *i.e*. hydroxyapatite column, was carried out using PBS buffer without chelator or dithiothreitol (DTT). Cystathionine γ-lyase (CSE) was expressed and purified following the described procedure[35]. hTrxR was overexpressed in BL21(DE3) cells and purified as follows. *E. coli* cells were resuspended in 100 mL extraction buffer (50 mM Tris-HCl, pH 7.5, 30 mM KCl, 5 mM DTT, 1 mM EDTA, 1 mM pMSF and 2 tablets of cOmplete™ protease inhibitor cocktail) and sonicated on ice for 5 min (10″ on; 40″ off). Following centrifugation for 60 min at 20,000 rpm and 4 °C, the supernatant fraction was submitted at first to a streptomycin sulfate (2.5 % w/v) precipitation step, then treated with pancreatic DNAse I (3 mg) after centrifugation, and lastly precipitated with saturated ammonium sulfate (30-85 %) for 60 min at 4°C. The solution was centrifuged as described above, and the yellow pellet was resuspended in 50 mM Tris-HCl, pH 7.5, 1 mM EDTA (buffer A) and dialysed overnight against the same buffer at 4°C. The protein extracts were then loaded on a HiPrep DEAE FF 16/10 column (Cytiva). hTrxR was eluted with a linear gradient from 10 to 500 m M KCl in buffer A. Fractions containing hTrxR, as judged by SDS-PAGE 12%, were pooled, concentrated with a centricon YM10, and applied onto a HiLoad Superdex 200 (Cytiva) equilibrated with buffer A complemented with 100 mM KCl. The enzyme was submitted to an isocratic elution and fractions containing pure hTrxR were dialyzed against 50 mM potassium phosphate, pH 7.5, 1 mM EDTA and stored frozen at −80 °C.

### 2.3. Reactions between DJ-1 and 2,2-dithiodipyridine

To a 180 μM solution of 2,2-dithiopyridine in the suitable buffer was added an 18 μM solution of DJ-1 or its mutant C106S in the same buffer and the absorbance at 343 nm recorded over time. Absorption-time profiles were fitted with SigmaPlot 10 to the double exponential A1[1 − exp(−*k*_obs1_ t)] + A2[1 − exp(−*k*_obs2_ t)] + B or the mono exponential A[1 − exp(−*k_obs_* t)] + B function.

### 2.4. Reactions between DJ-1 and sulfane sulfur donors

DJ-1 (12.5 μM) was incubated with various donors at room temperature (RT) for 30 or 60 minutes, and its esterase activity was determined by monitoring the slope of the absorbance at 405 nm *vs* time at 25 °C upon addition of 10 μL of the aforementioned solution to 190 μL of a 2.8 mM solution of *p*-nitrophenylacetate (*p*NPA, prepared in PBS without DTPA from a 200 mM stock solution in dimethylsulfoxyde (DMSO)). This activity was compared to the activity of a similar DJ-1 solution without donor (100%). Solutions of the donors without DJ-1 were also used as control and show no significant esterase activity unless otherwise stated. Kinetic data with Na_2_S_2_ were obtained at 25 °C as follows: to 5 mL of a 0.625 μM solution of DJ-1 were added 1.25, 2.50, 5.00 or 7.50 μL of a 10 mM solution of Na_2_S_2_ (final concentrations: 2.5, 5.0, 10.0 and 15.0 μM). The esterase activity was recorded at various time intervals by monitoring the slope of the absorbance at 405 nm *vs* time, upon addition of 1 mL of the aforementioned solution to 3 μL of a 200 mM solution of *p*NPA in DMSO. The pseudo-first order rate constants *k*_obs_ were obtained for each concentration by fitting the data with SigmaPlot 10 to the mono exponential function A[1 − exp(−*k*_obs_ t)] + B. The bimolecular rate constant *k*_inac_ is the slope of the linear fit obtained from the plot of *k*_obs_ *vs*[Na_2_S_2_].

DJ-1 protection of guanosine monophosphate (GMP) against modification induced by methylglyoxal (MGO) was determined by LC-MS. GMP (40 μM) was incubated with MGO (400 μM) and either wt DJ-1 (4 μM) or wt DJ-1 + Na_2_S_2_ (4 μM, from a stock solution prepared by incubating 40 μM DJ-1 with 60 μM Na_2_S_2_ for 20 min at 20°C) for 12 min at 37°C. Data were calculated by measuring the surface area of the glycated GMP peak (m/z = 436.09, ESI+) or lactate (m/z = 89.00, ESI-). 100% protection corresponds to the reaction [GMP + MGO + DJ-1] while 0% protection corresponds to the reaction [GMP + MGO], since 6 μM Na_2_S_2_ did not influence the reaction between GMP and MGO.

### 2.5. Reactivation of persulfidated DJ-1

A 13 μM solution of DJ-1 was inhibited by reaction with 15 μM of Na_2_S_2_, leading to an 80% drop of its esterase activity (see below). This solution was then incubated with either 1 mM DTT, the system hTrx/hTrxR/NADPH (8 μM/2.5 μM/400 μM) or the system hTrx/DTT (8 μM/1 mM) and the esterase activity recorded as described above at various time points.

### 2.6. Differential Scanning Fluorimetry experiments

Solutions of DJ-1 (50 μM) in PBS were incubated with/without 65 μM Na_2_S_2_ or 350 μM H_2_O_2_ for 20 minutes at RT, then desalted with Micro Bio-Spin 6 columns (Bio-Rad). To 22.5 μL of the resulting solutions was added KCl (final concentration 100 mM) and 2.5 μL of a 50x stock solution of Sypro orange in water. DSF assays were run using qPCR plates using the following parameters: samples were heated from 10 to 95 °C at a rate of 2 °C/min, and fluorescence recorded using the FRET channel. Noteworthy, DTT (1 mM) was added to the sulfinylated form of DJ-1 prior to qPCR as its omission precluded the recording of a clean melting curve. Data were plotted as the first derivative of fluorescence as a function of temperature, whose peak corresponds to the melting temperature (Tm). Identical Tm values were obtained by running the assay after mixing 20 μL of a 50 μM solution of DJ-1 in PBS without DTPA with 2.5 μL of a 650 μM solution of Na_2_S_2_ and 2.5 μL of a 50x stock solution of Sypro orange.

### 2.7. In gel detection of persulfidation

i. With purified proteins: wt or mutant (C106S) DJ-1 (approx. 1.5 mg/mL in PBS, 75 μM) were first treated with/without sodium disulfide (100 or 150 μM final concentration) for 20 minutes at RT, then sodium dodecyl sulfate (SDS) was added to reach a 2% final concentration. The resulting mixtures were then treated at 37°C with 20 mM NBD-Cl for 1 hour. Proteins were then precipitated with water/methanol/chloroform (4/4/1) and the resulting pellet was re-suspended in Hepes buffer (50 mM, pH = 7.4) containing 2% SDS, and further incubated at 37°C with Daz-2:Cy5 mix (50 μM) for 30 minutes [30]. After precipitation and resuspension of the pellet as above, the solutions were submitted to a denaturing non-reducing 12% criterion XT Bis-Tris gel (Bio-Rad).
ii. With *E. coli* extracts: The plasmids for wt or mutant C106S DJ-1 were transformed into BL21(DE3) *E. coli* strain and the bacteria were grown overnight in LB medium containing 100 μg/mL ampicillin. Next, the overnight culture was added at 2% v/v to LB medium supplemented with ampicillin (100 μg/mL). Bacteria were grown at 37°C until OD600 = 0.6, and IPTG (0.2 mM final) was added to induce protein expression. If needed, thiosulfate (25 mM) or cysteine trisulfide (1 mM) [36] was added 30 minutes after induction. After 5 h culture at 37°C under mild shaking (150 rpm), bacteria were pelleted, resuspended in PBS buffer supplemented with cOmplete^™^ protease inhibitor cocktail (Roche) and 4-Chloro-7-nitro-2,1,3-benzoxadiazole (NBD-Cl, 25 mM) and sonicated twice (20″ on, 60″ off) on ice. After 1 hour at 37°C, the resulting extracts were centrifuged at 20,000 rpm for 20 minutes, then the proteins precipitated using methanol and chloroform and further treated as previously described [30], before loading on a denaturing non-reducing 12% criterion XT Bis-Tris gel (Bio-Rad).

### 2.8. Molecular dynamics

The crystal structure of the DJ-1 dimer was downloaded from the Protein Data Bank (PDB ID: 3SF8, https://www.pdb.org [19b]). The dimer formed by subunits A and B was prepared using the Prepare Protein module in Biovia Discovery Studio_®_ (DS) 2021 with the default parameters and the CHARMM force field. Briefly, all crystallographic water molecules were removed. Bond orders were assigned, hydrogen and missing atoms were added. The protonation states on protein were adjusted at pH 7.4. C106 was deprotonated and E18 protonated on the basis of previous studies, [19] sulfur or oxygen atoms were added to the sulfonate form of C106 when required, and the structures were minimized.

To assess the influence of persulfidation at C106 and compare it to the wt or sulfinylated protein, 70 ns molecular dynamics (MD) simulations were run using the CHARMM36m force field and the NAMD protocol [37] implemented in DS 2021. Proteins were solvated in a cubic box using a TIP3P water model. Periodic boundary conditions were applied with a minimum distance of 10 Å from periodic boundary and Na+Cl-counter ions were added to neutralize the system. Solvated complexes were subjected to 2 cycles of energy minimization (1000 steps of Steepest Descent algorithm then 20000 steps of ABNR algorithm) followed by 500 ps of heating from 50 to 300K at constant volume, 1 ns of equilibration at 300K and 50 ps of production in the NPT ensemble (300K, 1 atm). All MD simulations were performed under NPT conditions (300K, 1 atm). Langevin Dynamics- and Langevin Piston methods were applied to control the tempera-ture and the pressure. Short-range electrostatic and Van der Waals interactions were computed with a 12 Å cutoff distance and long-range electrostatic interactions were treated by the Particle Mesh Ewald (PME) method. All bonds with hydrogen atoms were held rigid using the SETTLE algorithm. RMSD and RMSF values, as well as distance and interface energies were calculated using the Analysis Trajectory tool of DS. Electrostatic potentials were calculated using the CHARMm PBEQ module implemented in DS.

## 3. Results

As mentioned above, DJ-1 is a target for many PTMs including persulfidation. Several reaction pathways may result in the persulfidation of a cysteine residue under biological conditions [38]. We however focused at first on the reaction of hydrogen sulfide (H_2_S) with an oxidized cysteine residue of DJ-1 and then on the reaction of a protein cysteine residue with sulfane sulfur [33] donors to access a persulfidated form of DJ-1. We carried out our reactions in a buffer containing a chelator for metal ions (DTPA), because DJ-1 has been proposed to bind copper [39,40] and zinc [41] *in vitro*, even if these hypotheses were recently ruled out in a cellular context [42].

### 3.1 C106 of recombinant human DJ-1 is the most thiophilic cysteine

To obtain persulfidated DJ-1, we first envisioned the method proposed by Pan *et al*. [43], based on the formation of a reactive S-S bond and its subsequent reduction by sodium hydrosulfide. Because DJ-1 possesses three cysteines, we turn our attention to 2,2’-dithiopyridine (DTP) to generate the mixed reactive disulfide thiopyridine(TP)-C106 DJ-1. Indeed, DTP should allow the selective labelling of the low-p*K_a_* C106 as it reacts with thiols (Scheme 1) even at low pH [44].

**Scheme 1.**
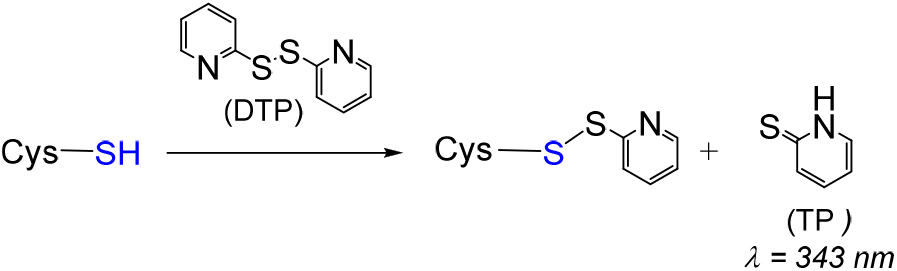
General reaction between cysteine and 2,2’-dithiopyridine (DTP).

The release of thiopyridine TP upon reaction between DJ-1 and DTP can easily be monitored by UV-visible spectroscopy at 343 nm. The kinetic data obtained at pH = 7.4 (Figure 1, blue data) can be nicely fitted with a double exponential function, yielding *k_obs_* values of 1.667 ± 0.002 and 0.088 ± 0.004 min^−1^. This observation suggests either the reaction of two of the three Cys residues of DJ-1 with DTP, or the reaction of a single Cys residue with DTP followed by reduction of the mixed reactive disulfide by a second Cys residue. To distinguish between these two processes, we performed a similar experiment with the mutant C106S DJ-1. Its reactivity with DTP yields a single *k_obs_* (0.098 ± 0.006 min^−1^) similar to the slower *k_obs_* previously measured with the wt protein, ruling out the second hypothesis. Thus, two cysteine residues of DJ-1 react with DTP, and C106 is the most thiophilic one. As expected, the selectivity for C106 is somewhat increased at pH = 6.0 for which the reaction of C46/C53 is significantly slower (*k_obs_*=0.034 ± 0.012 min^−1^) while the reaction of C106 is marginally faster (*k_obs_*=1.976 ± 0.238 min^−1^).

Next, to obtain persulfidated DJ-1, we removed excess DTP and incubated the mixed reactive disulfide TP-DJ-1 with a 20-fold excess of hydrosulfide at 25°C for 30 min. Unfortunately, we did not observe the expected release of TP from the mixed disulfide upon sodium hydrosulfide (NaSH) treatment, indicating that the generation of a persulfidated form of DJ-1 requires another approach.

### 3.2. Recombinant human DJ-1 is inactivated by sulfane sulfur donors

We next turned our attention towards the reaction of DJ-1 with sulfane sulfur sources. To detect a potential modification of the protein properties, we took advantage of the esterase activity of DJ-1, which releases the chromophoric *p*-nitrophenoxide (λ = 405 nm) upon incubation with *p*-nitrophenyl acetate (*p*NPA) [18]. We started our investigation with the garlic-derived diallyltrisulfide (DATS), a natural source of reactive sulfur species (RSS)[45]. No loss of esterase activity was observed upon incubation of an excess (5-10 equiv.) of DATS with DJ-1, even after several hours. However, in the presence of an additional 1 mM of glutathione (GSH), the enzyme was dose- and time-dependently inhibited (Figure 2A), clearly advocating for the reaction between a cysteine residue of DJ-1 and (an) intermediate species formed by the reaction between GSH and DATS [46].

Amongst those [46,47], we first ruled out the implication of H_2_S, because it is unlikely on a mechanistic ground and because NaSH alone did not inhibit DJ-1. Interestingly, persulfides (R-SSH), generated either from the synthetic precursor P* developed in our group [48] or enzymatically produced by cystathionine γ-lyase (CSE) [49], led to a clear inhibition of DJ-1 (Figure 2B). However, glutathione persulfide, generated from GSH and phenylthiosulfonate [50], failed to inhibit DJ-1 suggesting that the steric hindrance of GSSH prevents its reactivity with a buried cysteine. Finally, the two polysulfides sodium salts di- and tetra-sulfide proved to be the most efficient donors to inhibit DJ-1. For instance, stoichiometric amount of sulfane sulfur with respect to DJ-1 (1 equiv. of Na_2_S_2_, or 0.33 equiv. of Na_2_S_4_) leads to ~ 80% loss of activity after 30 minutes and does not further inhibit DJ-1 after 60 min, advocating for a fast reaction rate between polysulfide sodium salts and the C106 of DJ-1.

To get further insight into DJ-1 inhibition provoked by sodium disulfide, we next carried out kinetic investigations using the approaches proposed by Mangel *et al*. [51] to study fast-acting proteinase inhibitors. First, we recorded the release of *p*-nitrophenoxide over time, in the presence of increasing amounts of sodium disulfide (Figure S1). As expected, a decreased plateau level was observed when the concentration of Na_2_S_2_ increased. Interestingly, for 2 to 20 μM concentrations, the rate of hydrolysis remained constant after 200 s and approximatively three time the rate recorded with the mutant C106S in absence of sodium disulfide, suggesting the existence of a residual activity for DJ-1 treated with Na_2_S_2_. However, we experienced difficulty to reproduce these experiments and extract a bimolecular rate constant for DJ-1 reactivity with sodium disulfide using this continuous activity assay, most probably because of side reactions involving sodium disulfide and the substrate *p*NPA. To overcome this problem, we pre-treated DJ-1 with 4 different concentrations of Na_2_S_2_ for varying periods of time before recording its esterase activity. Reproducible data were thus obtained, which are presented in Figure 3A. The slope of the straight lines obtained for each Na_2_S_2_ concentration gave pseudo first-order rate constants kobs that, once plotted against the various Na_2_S_2_ concentrations gave a bimolecular rate constant k_inac_ of (1.69 ± 0.10) × 10^3^ M^−1^.s^−1^ (Figure 3B). Noteworthy, a k_inac_ of 3.8 ± 0.3 M^−1^.s^−1^ was observed in a similar experiment performed with hydrogen peroxide. The later value compares well with the one previously reported by Andres-Mateos *et al*. (0.56 ± 0.05 M^−1^.s^−1^) obtained using a different assay [52].

Finally, we performed experiments to assess the potential impact of Na_2_S_2_ on the protective activity of DJ-1 against methylglyoxal-induded (MGO) modification of GMP. As expected for an impaired DJ-1 activity, less lactate is produced and more GMP-MGO adduct is detected when DJ-1 is treated with the polysufide (Figure 3C). We did not investigate the exact mechanism underlying this protection, as recent kinetic studies reassessed DJ-1 as a glyoxalase rather than a deglycase [10,53].

### 3.3. DJ-1 is slowly reactivated in vitro by DTT, but not by hTrx, hTrxR or GSH

Usually, modification of a protein by sulfane sulfur donors is reversible and the reversal is often catalyzed by the thioredoxin and glutathione systems [54,55]. Accordingly, we monitored the reactivation of inhibited DJ-1 by various reducing systems. Incubation with DTT restores the activity of the inhibited protein, albeit with a slow *k*_reac_ of 0.075 ± 0.005 M^−1^.s^−1^ (Figure 4A). However, glutathione, the mammalian system NADPH/hTrx/hTrxR or the stoichiometric system hTrx/DTT were unable to reactivate the esterase activity of DJ-1 in our hands, pointing to the need for a specific reducing system to reactivate DJ-1. Interestingly, pre-incubation of the inhibited enzyme with H_2_O_2_ before addition of DTT or of the hTrx system did not affect the reactivation. Additionally, the oxidation of DJ-1 with H_2_O_2_, in the presence or absence of equimolar concentration of H_2_S, was irreversible regardless of the reducing agent (Figure 4A).

### 3.4. The modification induced by sodium disulfide increases the thermal stability of DJ-1

Structural modifications induced on human DJ-1 by hydrogen peroxide are accompanied by a change in its melting temperature (Tm) [56], as determined by Differential Scanning Fluorimetry (DSF) which gives information on protein stability. Accordingly, inhibited DJ-1 by sulfane sulfur donors should also exhibit a variation in its Tm as compared to native DJ-1. We thus determined the Tm of both proteins in PBS using differential scanning fluorimetry (Thermofluor). Native DJ-1 has a Tm of 60.0 °C (Figure 4B), which is identical to the value previously reported for DJ-1 under similar experimental conditions [53]. The incubation of DJ-1 (50 μM) with sodium disulfide (65 μM) shifts this Tm to 65.5 °C, advocating that the modification of DJ-1 by sodium disulfide stabilizes the protein. Interestingly, we were under these conditions unable to record a clean melting curve for the oxidized form of DJ-1. However, in the presence of DTT, we were able to reproduce the Tm of 75°C previously reported [56]. The reducing agent is thus critical, and it also shift the Tm of the persulfidated form to 70°C, but leads to a broader melting curve. Both PTMs therefore thermally stabilize DJ-1, and sulfinylation is slightly more stabilizing.

### 3.5. DJ-1 is persulfidated in vitro by polysulfides at C106

To clearly identify the modification responsible for the drop of enzymatic activity upon DJ-1 treatment with sulfane sulfur donors, we used the selective “tag-switch” method developed by Filipovic [30]. Briefly, persulfides are blocked as an activated mixed-disulfide by reaction with 4-chloro-7-nitrobenzofurazan (NBDCl), and the disulfide bond is reduced by a fluorescent dimedone derivative, leading to a fluorescent protein conveniently detected by in-gel fluorescence. In the absence of Na_2_S_2_, a weak signal was detected by florescence (Figure 5A). However, incubation of DJ-1 with sodium disulfide leads to a strong fluorescent spot on the gel. Interestingly, weak fluorescence was detected when the mutant C106S was incubated with or without Na_2_S_2_, indicating that DJ-1 is almost exclusively persulfidated at C106. This selectivity for C106 is confirmed by our initial data using a different assay (Figure S1bis).

### 3.6. Recombinant DJ-1 is endogenously persulfidated in E. coli

Finally, we investigated whether recombinant DJ-1 is endogenously persulfidated when overexpressed in *E. coli*, and if its persulfidation level could be influenced by sulfur compounds. When DJ-1 is overexpressed under standard conditions, persulfidation is clearly detected by in-gel fluorescence assay [30] (Figure 5C). The addition of 25 mM sodium thiosulfate to the culture medium after induction not only increases DJ-1 but also the global proteins’ persulfidation levels (expressed as the Cy5/CBB signal ratio, which we found more accurate in our experiments than the Cy5/488 signal ratio proposed previously [30]), in contrast to the addition of 1 mM cysteine trisulfide that had the opposite effect. It must be noted that endogeneous persulfidation was also unexpectedly detected when overexpressing the mutant C106S in *E. coli* (Figure S1ter).

### 3.7. Persulfidation affects the subunit interface (but sulfinylation does not)

In the absence of X-Ray data, we decided to use molecular dynamics (MD) simulations to evaluate the impact of C106 persulfidation on the structure and the stability of the protein. Root Mean Square Deviation (RMSD) plots (Figure 6A) show a good convergence of the trajectories after 25 ns and up to 70 ns indicating a good stability for the two systems. The average Cα-RMSD per residue which gives indications on the dynamics of individual amino acids of the dimer is plotted in Figure 6B.

**Figure 6.**
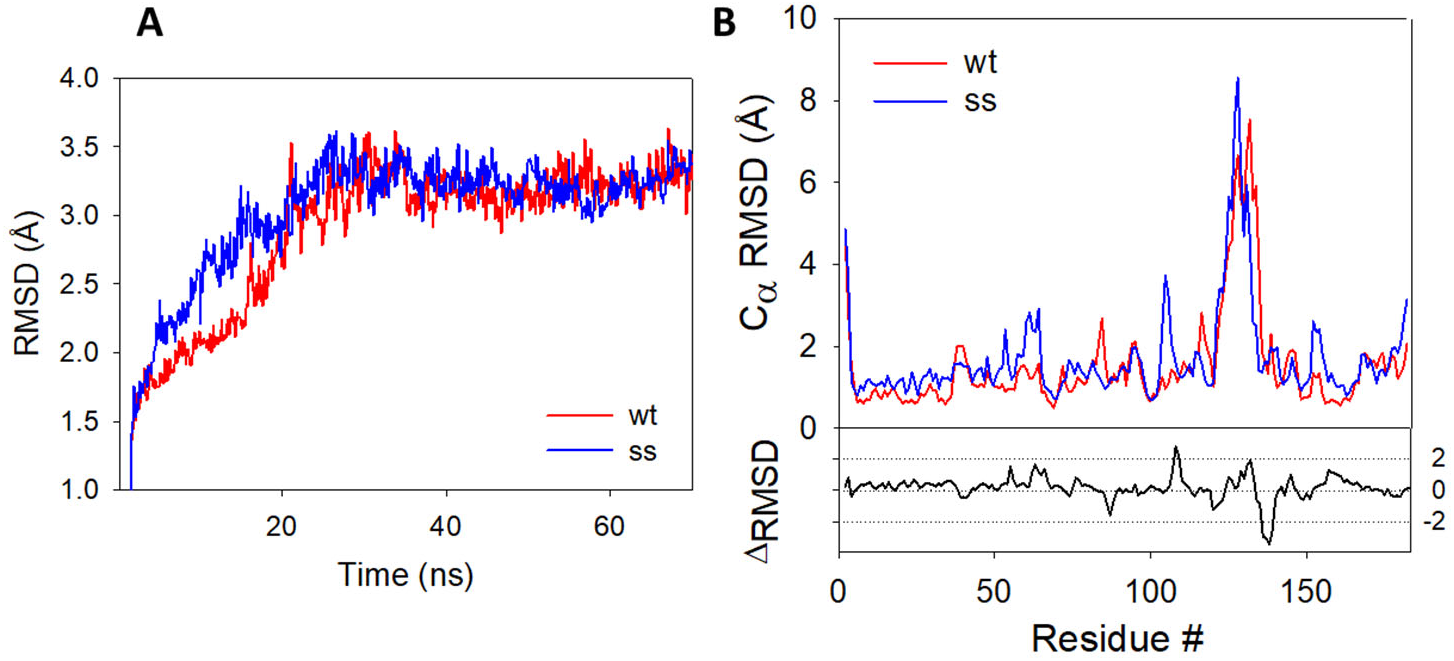
RMSD plots during 70 ns MD simulations. **(A)** RMSD plots obtained for the protein backbone of the wild-type (blue) and persulfidated (red) DJ-1 **(B)** Cα-RMSD of the amino acid residues from the wt (blue) and persulfidated (red) DJ1-1. RMSD plots represent the average of the RMSD value of each monomer.

The difference between the plot of the persulfidated and the wt proteins highlight several differences between the two trajectories. The most prominent deviations come from a partial loss of secondary structure on the loop connecting the β7 and α5 regions and containing the C106 (residues #106-109), and in the α6 region (residues #128-138) (Figure S2) in the persulfidated DJ-1. The addition of a sulfur atom to C106 just slightly impacts the main connections between this residue and its surrounding amino acid residues (G75, S155 and R156), with very similar distances recorded between these residues in both the wt and the persulfidated form (Figure S3). As previously proposed, in the B subunit of the wt, the deprotonated sulfur atom of C106 is stabilized [17] by H-bond interaction with protonated E18 and G75 (distance = 2.44 Å and 2.35 Å respectively). In subunit A, the sulfur atom does not interact with E18, a feature already reported by others [57]. Similarly, a weak interaction between E18 and the internal sulfur atom of the persulfide is noticed in the B subunit of the persulfidated form (distance = 3.96 Å). However, the outer sulfur atom of the persulfide remains mostly unstabilized and only interacts in a few conformations with the NH moiety of G75 (average distance of 3.78 Å). In addition, the latter is replaced in a few rare cases by an H-bond contact between G108 and the carbonyl group of C106 (Figure S3). The poor interaction network of the terminal sulfur may explain its higher fluctuation as shown by its higher Root Mean Square Fluctuation (RMSF) value compared to the one calculated for the inner sulfur atom of the persulfidated DJ-1 or the sulfur atom from C106 in the wt protein (1.53 *vs* 0.57 and 0.80 Å respectively) [58]. The same trend is followed for solvent accessibility, the additional sulfur atom being the most accessible one (SAS of 33.4, 7.5 and 11.8 Å^2^ for the additional S atom and the S atom of the persulfide, and the wt forms of DJ-1, respectively). Additionally, the persulfidation of C106 impacts the interface between the two subunits. Thus, crucial interactions implicated in the structural stabilization of the wt dimer [59,60] are either lost (R27A:R48B, G159A:L185B, Figures S4 & S5) or weakened (R28A:E15B, E18A:R28B and D49A:R27B, Figures S6 & S7) in persulfidated DJ-1 when compared to the wt. On another hand, a new interfacial H-bond contact appears between the guanidine group of R28 from the B subunit and the hydroxyl moiety of S47 from the A subunit (average distance of 3.15 Å *vs* 7.40 Å on the last 45 ns in the persulfidated *vs* wt, Figure S8). Overall, a significant decrease of the average interfacial interaction energy calculated on the last 20 ns is observed between persulfidated DJ-1 (−288.14 ± 16.87 kcal.mol^−1^) and the wt protein (−319.67 ± 12.89 kcal.mol^−1^). Finally, noticeable changes both in the hydrophobicity and electrostatic potentials are observed between the wt and the modified protein, in particular around the α6 helix that is the region showing the highest RMSD between the wt and the modified protein (Figure S9). Interestingly, C106 becomes more accessible after persulfidation, thus suggesting functions related to this exposure, *e.g*. intervention in transpersulfidation reactions or reversibility of the modification thanks to a dedicated depersulfidation system, while the SASA of the protein remain unchanged (14706 ± 175 *vs* 14979 ± 170 Å^2^).

Next, to compare persulfidation and sulfinylation, we generated the sulfinylated form of DJ-1 and performed MD simulations using the same procedure. A comparison between the final conformations of both oxidized proteins is presented in Figure S10. The structures of the persulfidated and sulfinylated DJ-1 are globally similar, but the latter does not show the partial loss of secondary structure observed in the α6 and α5-β7 regions of the former. Importantly, the active sites differ significantly, with strong interactions detected in the sulfinylated protein between the backbone of His126 and either the SO2 moiety (subunit A) or the carbonyl group of C106 (subunit B) (Figure S11). An additional H-bond contact between the sulfinic group and E18 additionally locks the conformation of C106 in subunit A, while G157 and H126 are in H-bond radius with the sulfinyl group in the B subunit. In addition, S155 strongly interacts with C106 backbone in the B subunit. Thus, the interactions observed in the persulfidated and wt forms between G75 and C106 are lost in the sulfinylated DJ-1. These results are in very good agreement with a previous study on oxidized and over-oxidized DJ-1 [57]. This new set of interactions results in a less mobile cysteine residue (the average RMSF of C106 is 0.98 Å for persulfidated DJ-1 and 0.53 Å for the sulfenylated form). Despite these significant changes in the active site, none of the interfacial changes observed upon persulfidation are detected in the sulfinylated form: the interaction R27A:R48B is even stronger than in the wt, while others important interactions G159:L185, D49:R27, R28:E15, S47:R28 remain intact (Figure S11). Additionally, various changes are observed at the protein surface (Figure S13).

## 4. Discussion

As the only thiol-containing residue of proteins, cysteine is a crucial player in redox sensing and signaling. For instance, the redox messenger hydrogen peroxide [61] and the gaseous transmitter nitric oxide [62] have been long known to signal at least partially through the modification of cysteine residues. More recently, hydrogen sulfide, a gaseous transmitter endogenously produced via enzymatic activity, has also been reported to signal through a post-translational modification of cysteine, first named S-sulfhydration[63] but more rigorously redenominated persulfidation[64]. This PTM, which converts a cysteine residue CysSH into the corresponding persulfide CysSSH, may result from the reaction between H_2_S and cysteine sulfenic acid (CysS-OH), the H_2_O_2_-oxidized form of cysteine[65]. In contrast, CysSSH may also be formed by the reaction between CysSH and bound sulfane sulfur species (BSS) [66]. BSS, which encompass sulfur-derivatives with a sulfur formal oxidation state of −I and 0 [66], are notably produced in cells by hydrogen sulfide [67] or 3-mercaptopyruvic acid [68] metabolism. Although their exact speciation is challenging to establish, their global concentrations (from high nM to 50-100 μM depending on the studies) [49,66] are orders of magnitude higher than those detected for hydrogen sulfide or H_2_O_2_ in various biological media under physiological conditions [66].

The outcome of persulfidation of cysteine residues on the function of targeted proteins is quite diverse [69,70] but sometimes conflicting. For instance, the first protein reported to be persulfidated (GAPDH) has been proposed to be activated or inhibited by this PTM [63,71]. Persulfidation may also induce intracellular relocation of targeted proteins, as observed with persulfidated GAPDH that is redistributed into the nucleus, enabling it to participate in H_2_S-mediated activation of autophagy [72]. Finally, this modification also regulates protein-protein interactions, as detected for the Keap1/Nrf2 system during the activation of the antioxidant response [73]. In addition to its role in intracellular signaling cascades, persulfidation has also recently been proposed as a protective mechanism against irreversible cysteine overoxidation during oxidative stress. Hydrogen sulfide may quench reactive sulfenic acid intermediates, thus preventing their further oxidation into sulfinic/sulfonic acids, and allowing the resulting persulfide (or persuf(e,i,o)nic species) to be reduced back to the thiol status by glutathione or the thioredoxin system [54,55].

DJ-1 has been known to be sulfinylated for years, with both C106 [22,23] and C46 [26,74] being target cysteines. The sulfinylation of C106 acts as a redox switch for DJ-1 activity, while the role of the modification of C46 is still obscure but likely of physiological significance since C46-SO**2**H has been described to be a substrate for sulfiredoxin Srx. More recently, DJ-1 has also been shown to experience persulfidation in mammalian cell lines [30] but neither the involved cysteine(s) nor the implications of this PTM on DJ-1 structure or activity have been analyzed. Accordingly, we aimed at elucidating the exact nature and consequences of the persulfidation process on DJ-1. At first, we expressed human DJ-1 in *E. coli*, and confirmed that it is partially persulfidated. The persulfidation level of DJ-1 increased in the presence of thiosulfate, a sulfur source for *E. coli*, but reduced in the presence of cysteine trisulfide, which has recently been proposed to be metabolized by *E. coli* into cysteine hydropersulfide [36]. Therefore, we expected to detect higher persulfidation yield of DJ-1 with cysteine trisulfide. However, a recent report confirms that cysteine trisulfides acts as an oxidative species leading principally to the oxidation of cysteine residues into mixed di-or trisulfides [75], which would account for the observed decrease in persulfidation levels when using this sulfane source. Next, the persulfidated form of DJ-1 was obtained by reacting purified wt DJ-1 with various sulfane sulfur donors, since the reduction of the activated disulfide bond formed from C106 and DTP by sodium hydrosulfide [43] had failed. Donors included cysteine hydropersulfide (enzymatically produced from cystine by CSE) or the polysulfide Na_2_S_2_ and Na**2**S4, used at physiologically relevant concentrations. Interestingly, glutathione hydropersulfide (formed *in situ* from glutathione and a chemical sulfur donor) [50] did not react with wt DJ-1, indicating that the size and/or charge of the sulfane sulfur donor govern the access to the reactive cysteine(s). Under these conditions, DJ-1 is selectively persulfidated at C106, as confirmed by the weak reactivity of the C106S mutant with sodium disulfide. This agrees well with our observations indicating that C106 is the most thiophilic Cys residue of DJ-1 when reacted with 2,2’-dithiopyridine (DTP) and with a previous study showing that glutathionylating agents mainly modify C106 [76]. This however contrasts with our observations suggesting that the mutant protein is also endogenously persulfidated in *E. coli*, even if the persulfidation of endogenous YajL, a member from the DJ-1 superfamily from *E. coli* [6] may account for this observation.

The post-translational oxidation of C106 by polysulfides inhibits DJ-1 C106-based activities (esterase or deglycase/glyoxalase activities). However, contrary to the sulfinylation that irreversibly inhibits the enzyme, the persulfidation is slowly reversible. Furthermore, in contrast to persulfidated PTP1B [77],HAS [54] or BSA [55] which are reactivated by the Trx and/or Grx systems, the persulfide of DJ-1 is solely reduced by DTT. This hints at a poor accessibility or electrophilicity of the inner sulfenyl sulfur of persulfidated C106. This would not only explain the lack of reactivity of H_2_S with the reactive mixed-disulfide form between C106 and DTP but also the absence of reactivity of the sulfenic acid of DJ-1 towards hydrogen sulfide (see below). This hypothesis, in agreement with the trans-nitrosylation (rather than the formation of a disulfide bond) observed from DJ-1 to the phosphatase PTEN [28], is also supported by MD simulation showing a larger solvent accessibility of the terminal sulfur atom.

To get insights into the kinetics of the sulfur transfer during the persulfidation of DJ-1 by small molecules we focused on the reaction between DJ-1 and sodium disulfide since the generation of the reactive persulfide from the systems CSE/cystine, P* or DATS/GSH are slow and limited by the rate of formation of the donor. We thus determined a rate constant of (1.69 ± 0.10) × 10^3^ M^−1^.s^−1^, which is, to our knowledge, the first reported for the reaction between a cysteine residue and a polysulfide. It is orders of magnitude higher than the bimolecular rate constant determined for the reaction of DJ-1 with hydrogen peroxide. Indeed, despite the low p*K_a_* of C106, its oxidation by hydrogen peroxide is slow (k = 0.56 M^−1^.s^−1^ [15] or 3.8 M^−1^.s^−1^ in this study, at pH 7.4) but in the range of those reported for the low molecular weight cellular thiols glutathione, cysteine or hydrogen sulfide (k = 0.9, 2.9 and 0.73 - 15 M^−1^.s^−1^ at pH 7.4, respectively) [78–80], or for proteins like human serum albumin (HSA) (k = 2.7 M^−1^.s^−1^) [81]. Additionally, DJ-1 does not stabilize the sulfenic form that quickly oxidizes to the sulfinic form[23].

**Scheme 2.**
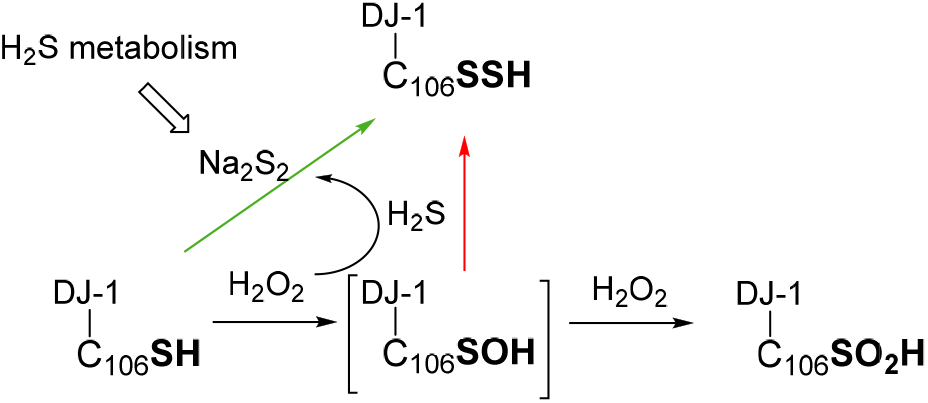
Simplified possible pathways for the formation of persulfidated DJ-1.

Therefore, contrary to other proteins in which the sulfenic acid is stabilized and may be quenched by hydrogen sulfide [65,82], the persulfidation of DJ-1 *via* the formation of its sulfenic acid (Scheme 2, red arrow) is in our opinion less likely than its reaction with polysulfides (Scheme 2, green arrow). This view is supported by our experiment showing that DJ-1 is fully and irreversibly inhibited in the presence of equimolar concentrations of H_2_O_2_ and H_2_S.

Finally, because C106 is the target of both sulfinylation and persulfidation, we next used additional techniques to investigate the differences induced by these two PTMs. The persulfidation of DJ-1 slightly stabilized the protein thermally compared to the wt form, as indicated by the 5°C variation of their respective Tm. Unfortunately, we were unable to obtain a clean melting curve with the sulfinylated form of DJ-1 and to directly compare the inherent stabilization afforded by each PTM, most probably because we had to work in the absence of DTT which reduces intra-disulfide bridges formed upon oxidation of DJ-1 [56,74]. Indeed, the determination of the Tm of various forms of DJ-1 is highly sensitive to the experimental conditions. For instance, totally different melting curves have been reported for the over-oxidized form of DJ-1 [57,83]). Our result would nevertheless suggest that the intrinsic stability of DJ-1 oxidized by Na_2_S_2_ or H_2_O_2_ differs and that their tertiary structures are dissimilar, which is supported by the lack of detection of persulfidated DJ-1 by the antibody directed against DJ-1 harboring a sulfinate. However, our MD studies indicate that both PTMs lead to tridimensional structures similar to the wt, a result already reported for sulfinylated and sulfonylated DJ-1 [57,84]. Nevertheless, at the local level, persulfidation results in a partial loss of the secondary structure and a decrease of the interfacial interaction energy similar to those observed in the pathological mutants like A104T [59], but absent in the sulfinylated form.

## 5. Conclusions

In conclusion, in this work we confirmed that DJ-1 is persulfidated not only in mammalian cells but also in *E. coli*. This PTM implicates cysteine C106, and on the basis of kinetic studies we propose that this oxidation takes place by the reaction of C106 with sulfane sulfur donors rather than by the reaction of its sulfenic form with hydrogen sulfide. Like sulfinylation, persulfidation inhibits two C106-based activities but the activity of the latter may be recovered in the presence of a reductant, albeit slowly. Additionally, various data suggest a structural difference between these two PTMs, which could both play a dedicated role in DJ-1 signaling or protective pathways or protein-protein interactions.

## Supplementary Materials

The following supporting information can be found at the end of the manuscript. Figure S1–S13: see the main text for description.

## Author Contributions

G. P. ran the preliminary experiments with sodium polysufides. E. G. conceived the study and performed the key experiments. N. M. produced and purified the wt and mutant DJ-1, ran the glycation tests and provided experimental support. L. LC. performed the molecular dynamics simulations. D. P. produced and purified hCSE and hTrxR and helped with data analysis. D. P. and J. D. provided intellectual input. E. G. wrote the manuscript with the help of all coauthors.

## Institutional Review Board Statement

Not applicable.

## Data Availability Statement

Raw gel images are available.

## Acknowledgments

We thank Booma Ramassamy for technical support and the Molecular Modelling Platform of UMR8601 - Université Paris Cité.

---

**Figure S1.**
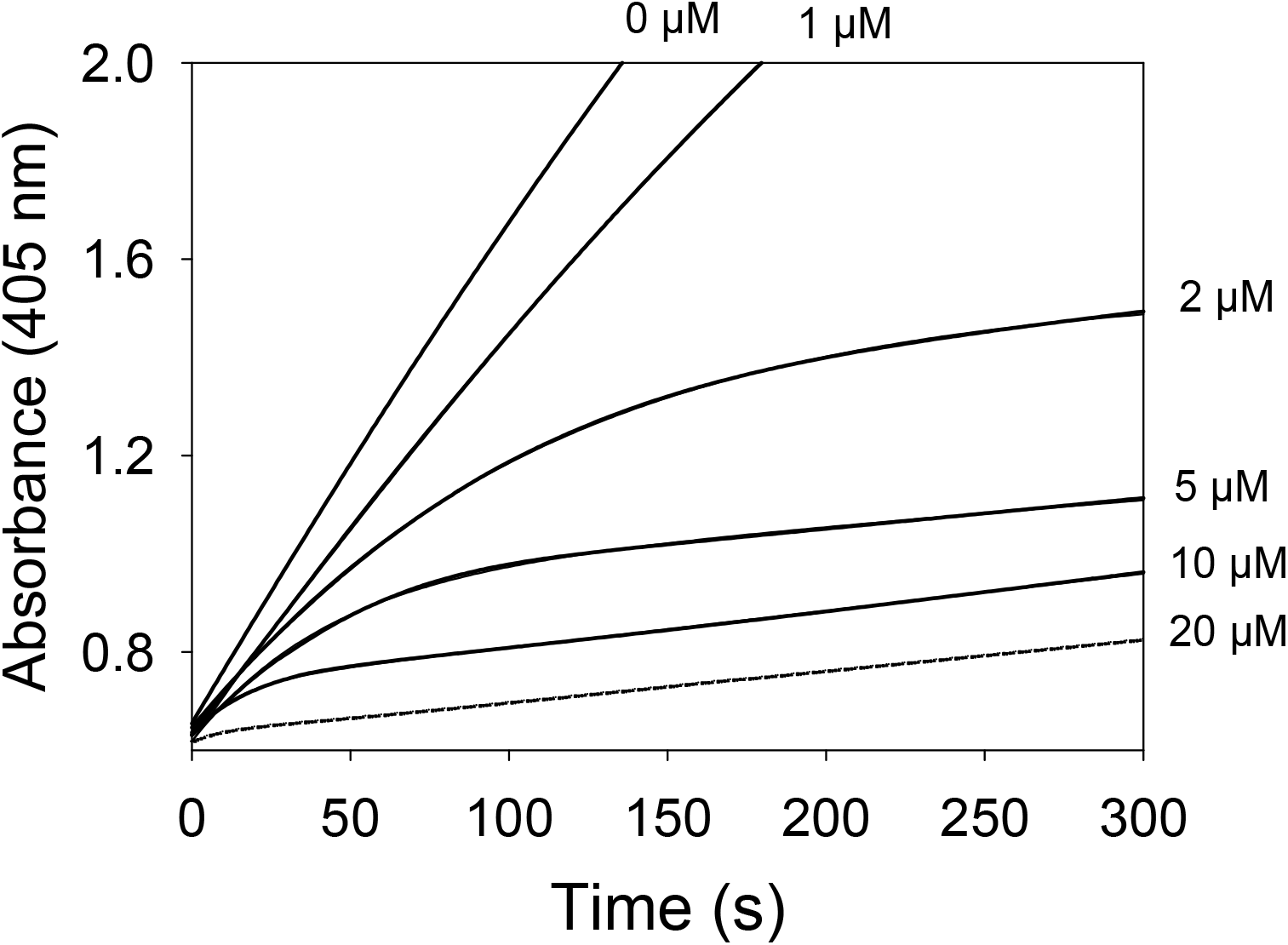
Kinetics of DJ-1 inactivation by sodium disulfide. Absorbance at 405 nm corresponding to the phenoxide released upon hydrolysis of pNPA (see the experimental section for conditions) recorded over time at 25°C after addition of DJ-1 (625 nM) to a solution of pNPa (2.8 mM) containing various concentrations of Na_2_S_2_.

**Figure S1bis.**
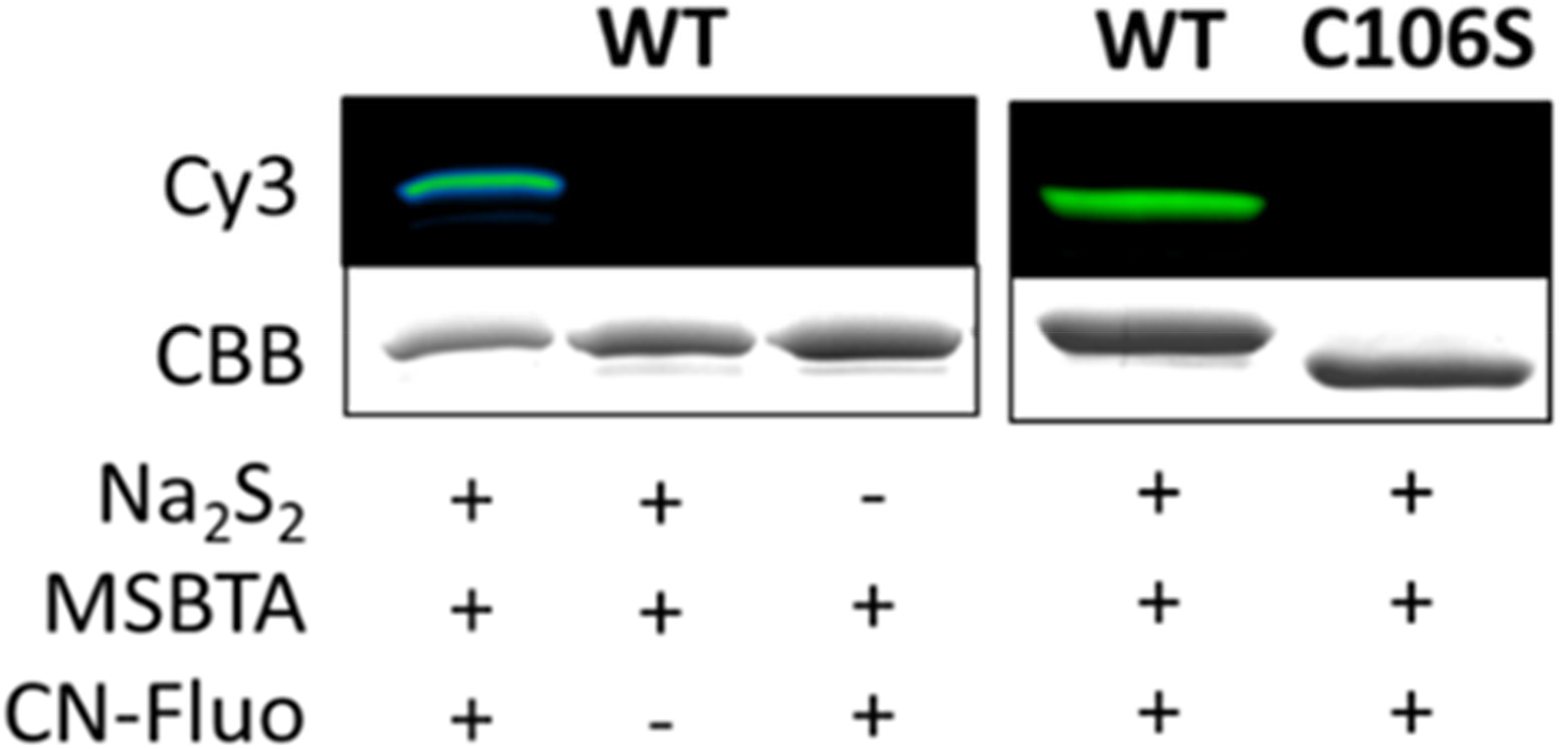
Detection of persulfidated DJ-1. In-gel fluorescence (Cy3-canal) and Coomassie Brilliant Blue staining scans obtained for WT or mutant C106S using the protocol from [1]

**Figure S1ter.**
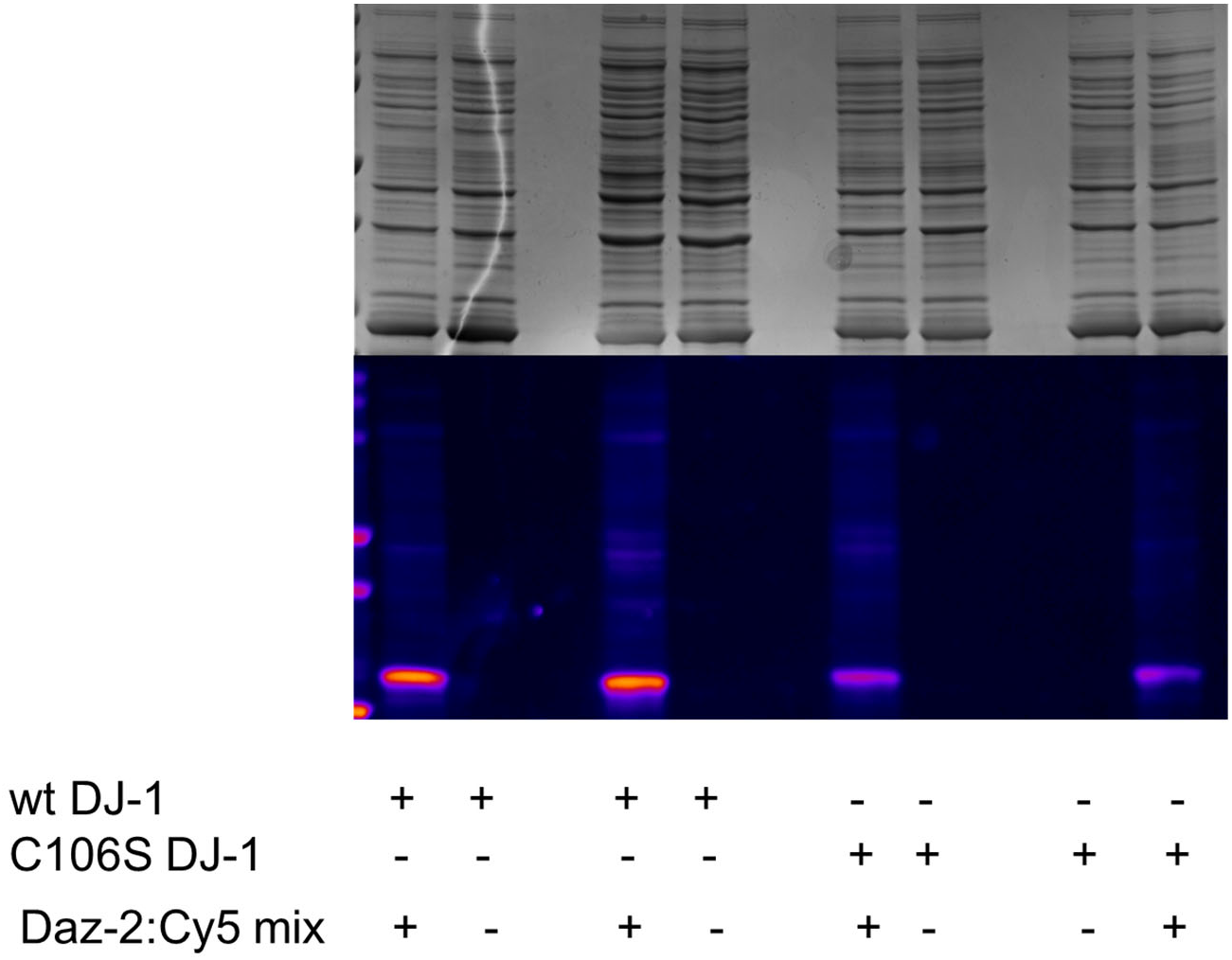
Persulfidation of C106S *vs* wt −DJ-1 in *E. coli*. In-gel fluorescence (Cy3-canal) and Coomassie Brilliant Blue staining scans obtained from *E. coli* overexpressing106S or wt DJ-1.

**Figure S2.**
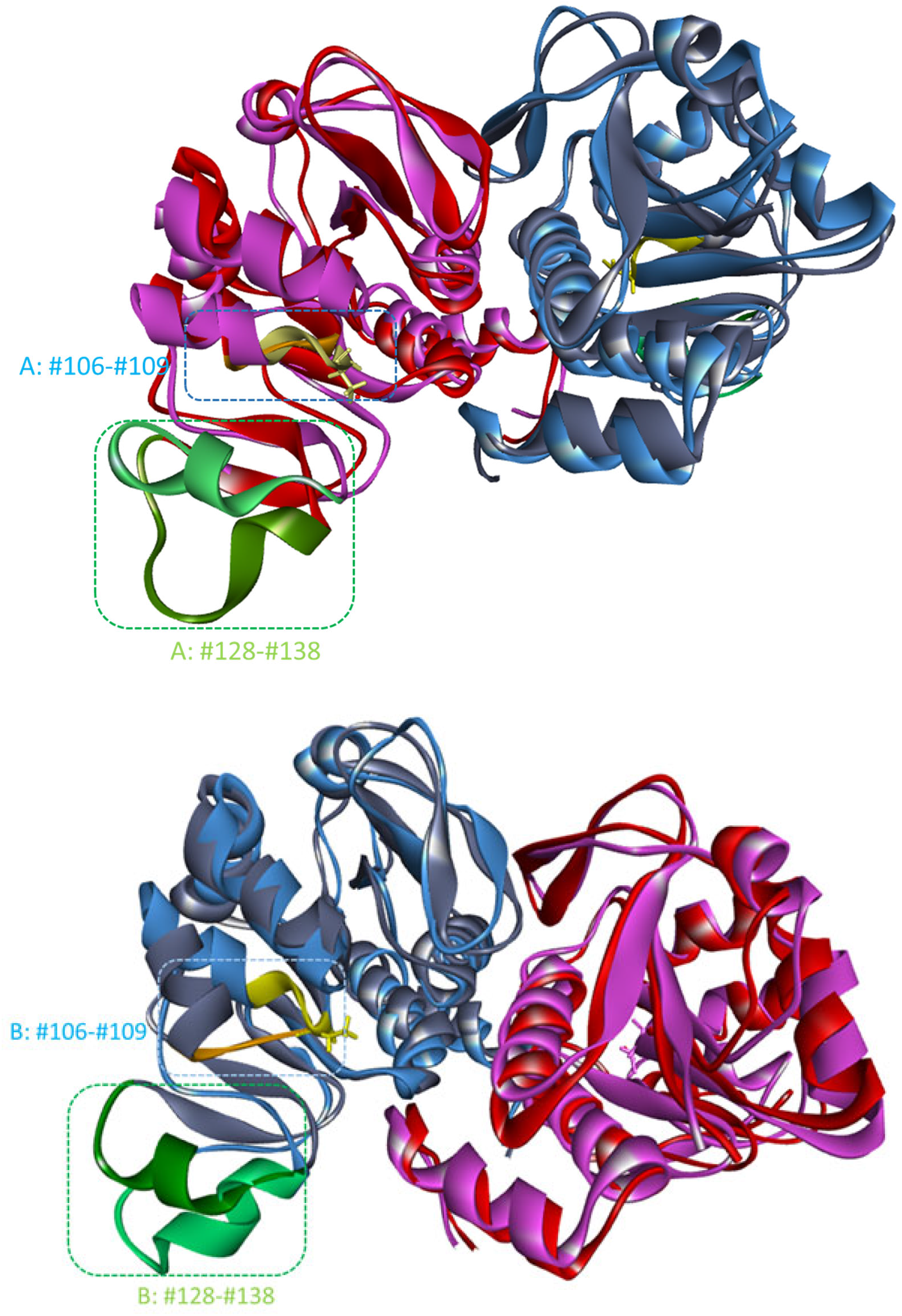
Final conformations of wt and persulfidated DJ-1. Structural comparison of representative snapshots of wt (pink and light blue) and persulfidated (red and dark blue) DJ-1. The two main regions showing important differences in RMSD are highlighted in yellow (wt) and orange (persulfidated) for residues #106-#109, and green (light for the wt, dark for the persulfidated form) for residues #128-#138.

**Figure S3.**
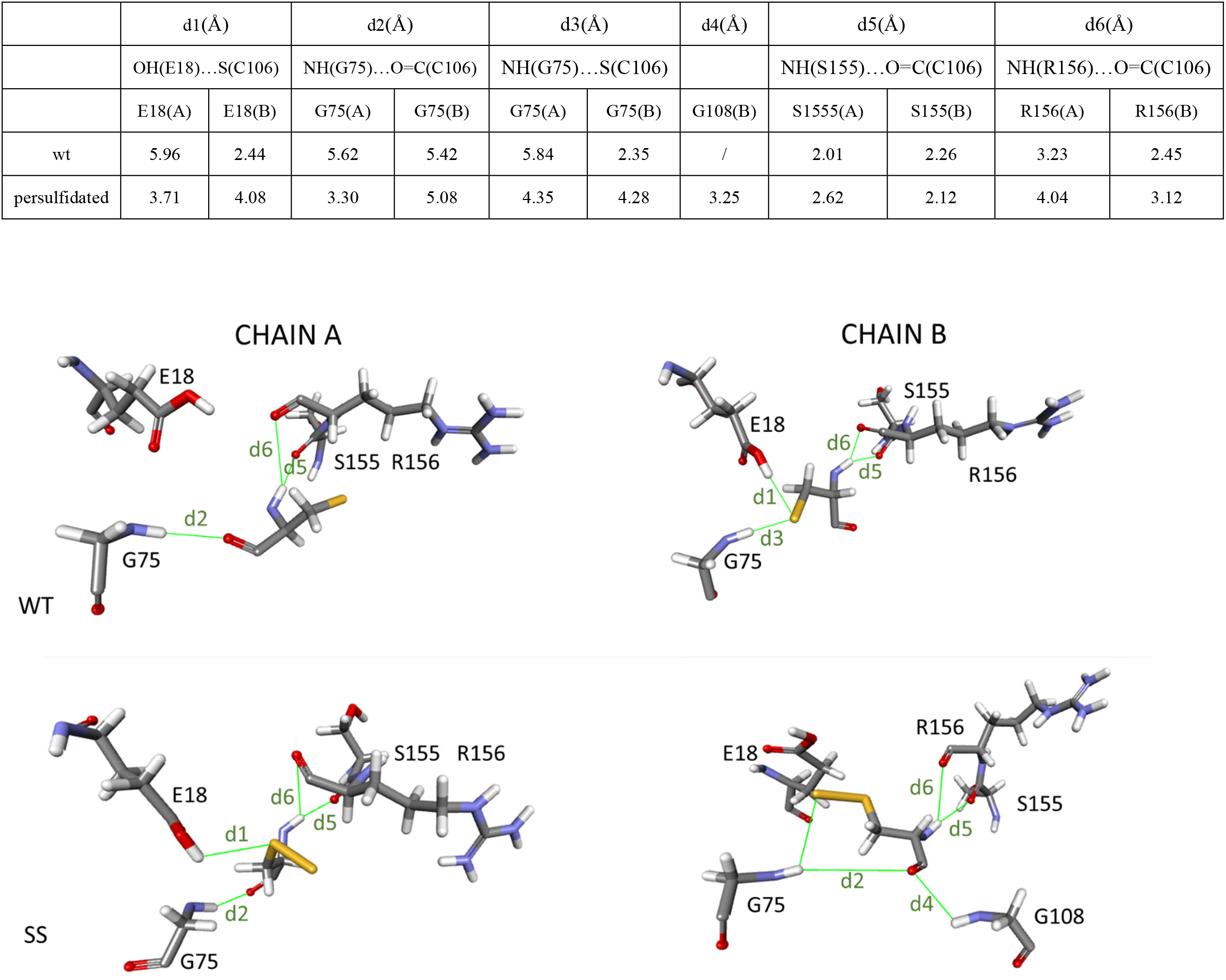
The main interaction between C106 and its neighbors are conserved upon persulfidation. Average distances (Å) recorded over the last 45 ns of the dynamics between C106 and interacting residues. Views of the amino-acids interacting with C106.

**Figure S4.**
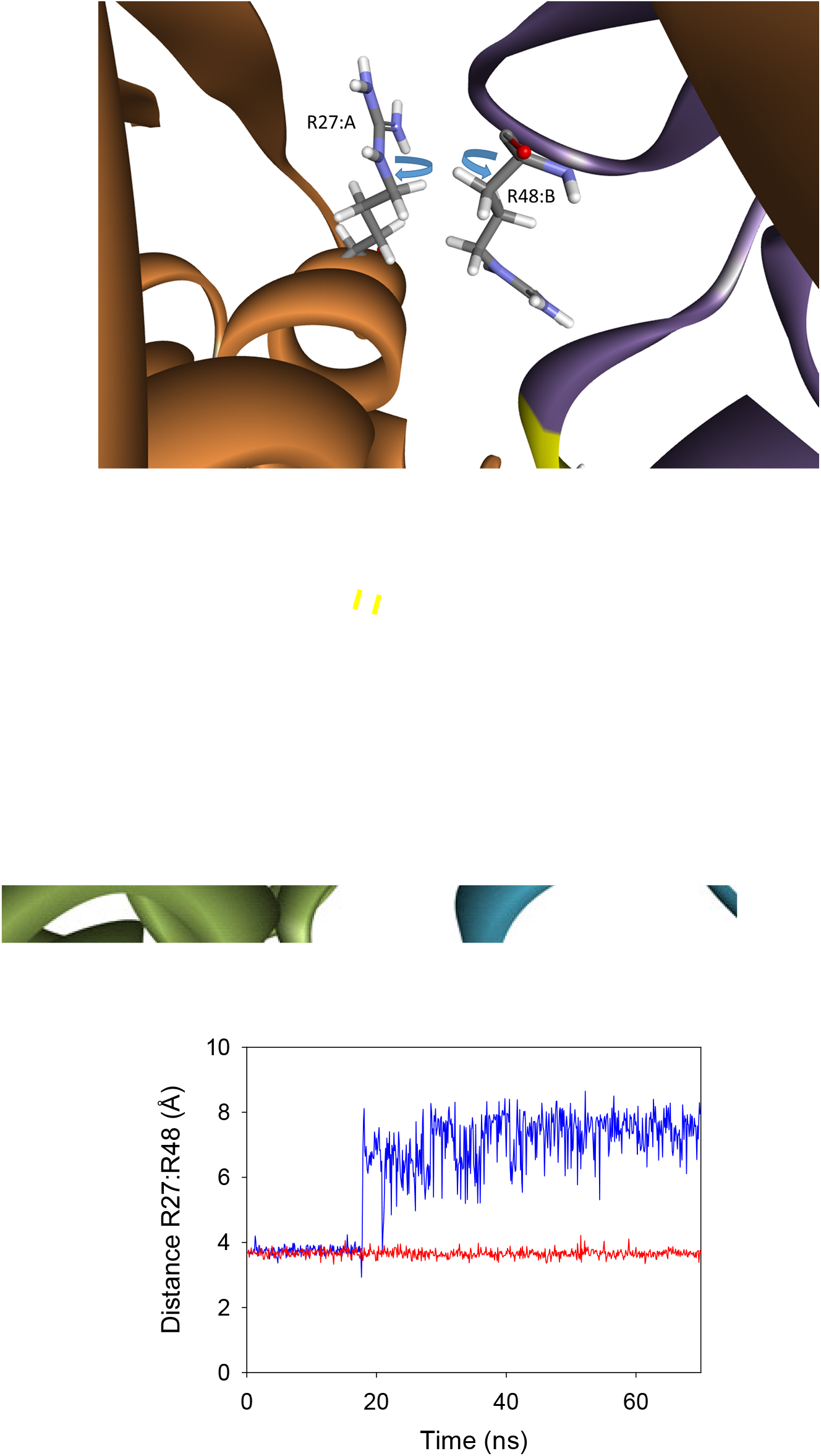
The interfacial H-bond intercation between R27:A and R48:B is lost after persulfidation. **A)** Representative snapshots obtained after 70 ns MD simulations of the wt (up) and persulfidated DJ-1 highlighting the loss of the interfacial H-bond interaction between R27 side chain (subunit A) and R48 backbone(subunit B), B) Time evolution of the distance between the centroid of the guanidine group of R27 and the carbonyle group of R48 over the course of the simulation in the wt (blue) and persulfidated (red) protein (bottom).

**Figure S5.**
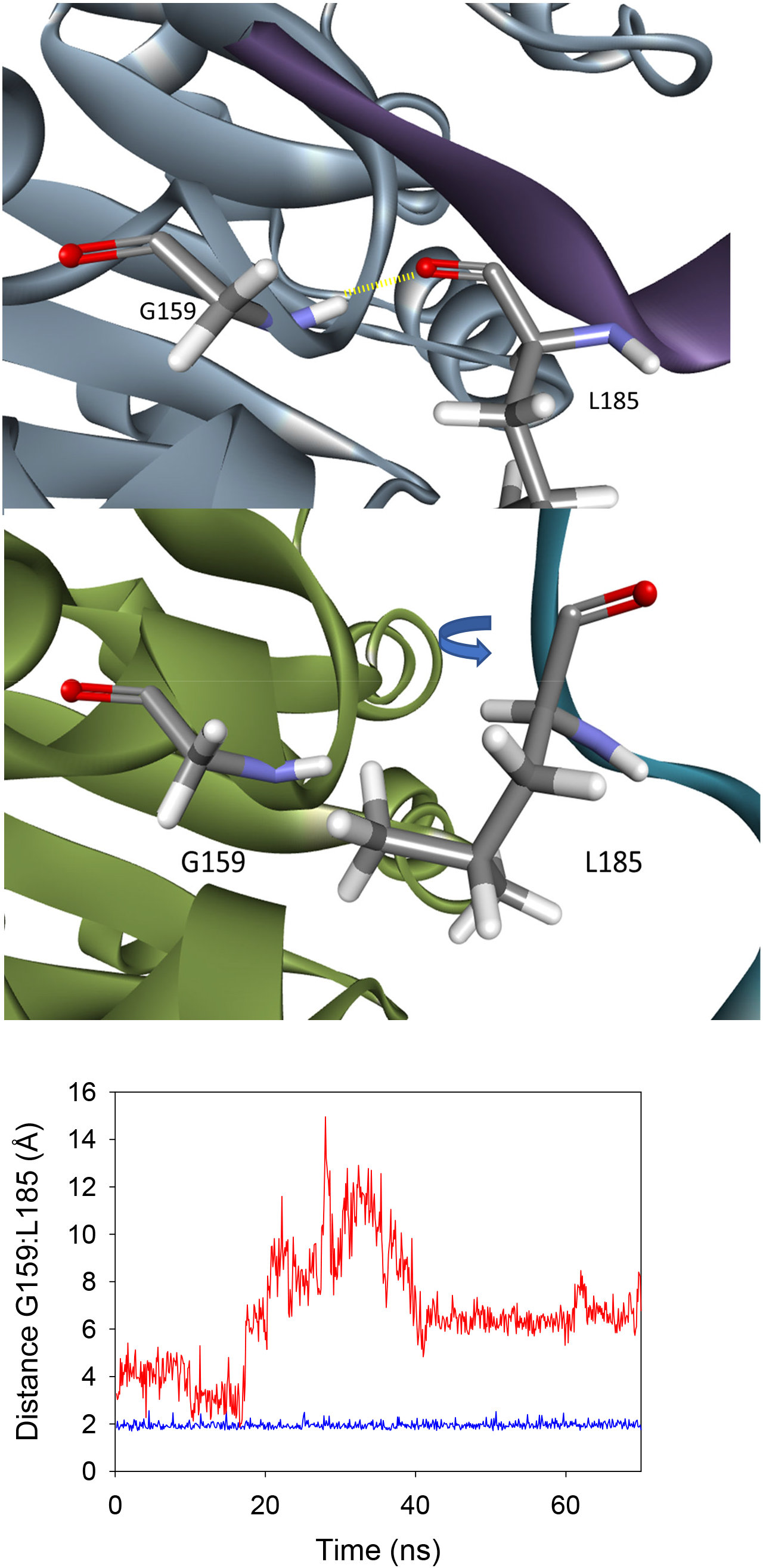
The interfacial H-bond interaction between G159:A and L185:B is lost after persulfidation. **A)** Representative snapshots obtained after70 ns MD simulations of the wt (up) and persulfidated DJ-1 highlighting the loss of the interfacial H-bond interaction between G159 and L185, B) Time evolution of the distance between the N-H group of G159 (subunit A) and the carbonyle group of L185 (subunit B) over the course of the simulation in the wt (blue) and persulfidated (red) protein.

**Figure S6.**
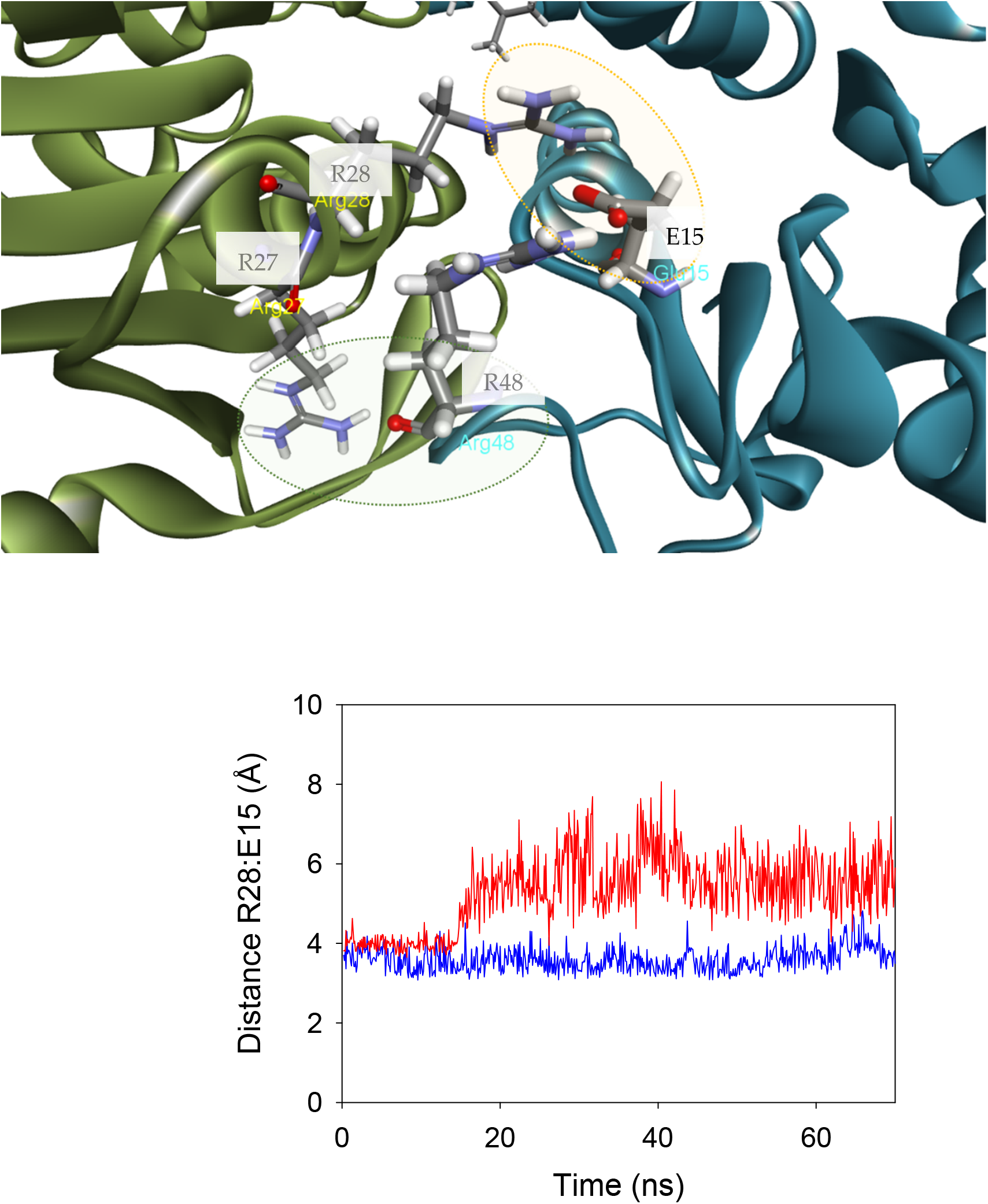
The interfacial interaction between R28:A and E15:B is weakened after persulfidation. **A)** Representative snapshot obtained after 70 ns MD simulations of the interfacial interaction between R28 and E15 (circled in yellow). B) Time evolution of the distance between the centroid of the guanidine group of R28 and the centroid of the carboxylate of E15 over the course of the simulation in the wt (blue) and persulfidated (red) protein.

**Figure S7.**
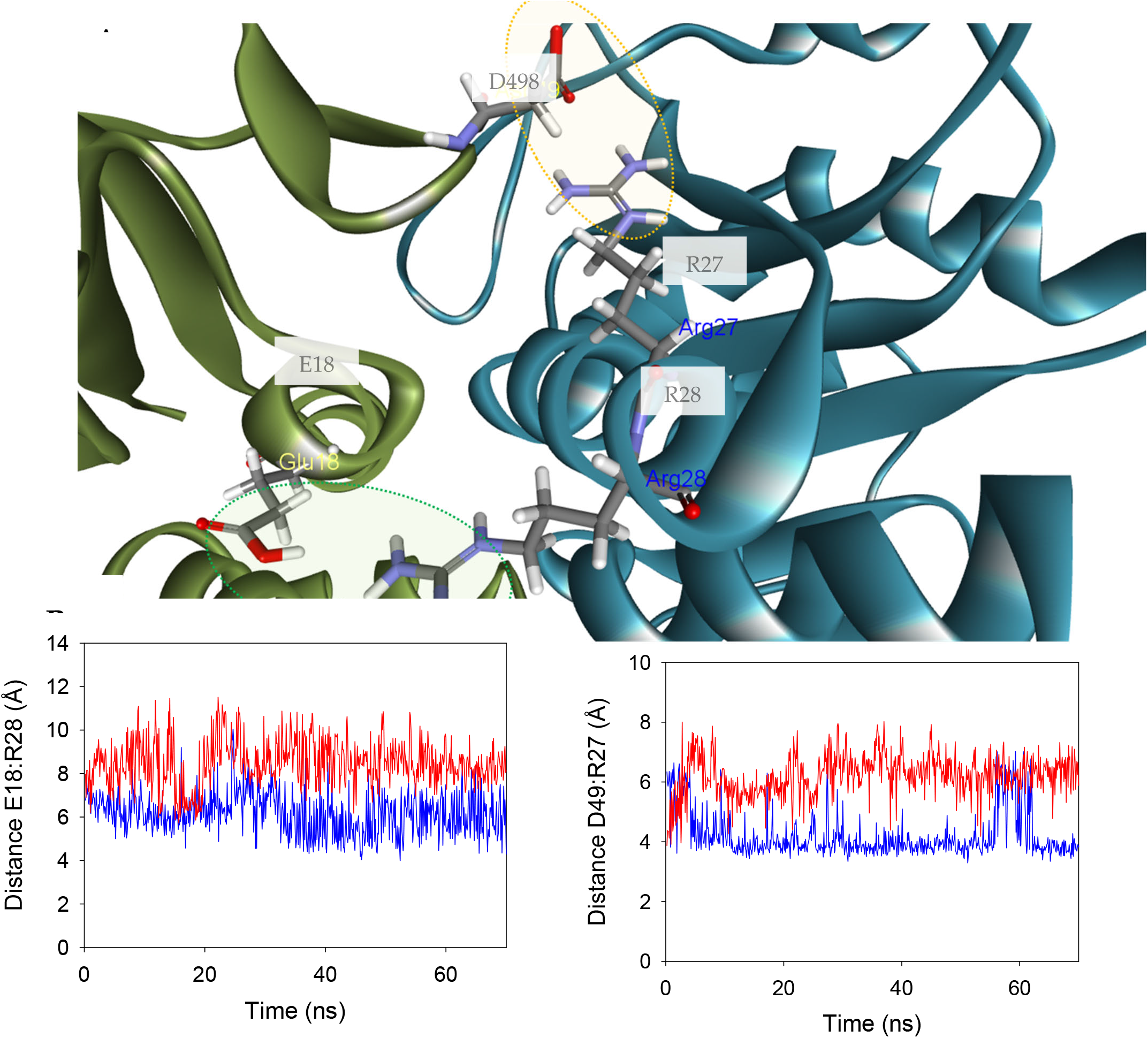
The interfacial interactions between E18:A and R28:B and between D49:A and R27:B are weakened after persulfidation. **A)** Representation of the interfacial interaction between E18 and R28 and between D49 and R27, B) Time evolution of the distance between the OH group of D18 and the centroid of the guanidine group of R28 (left) or between the centroid of the carboxylate groups of D49 and the centroid of the guanidine group of R27 (right) over the course of the simulation in the wt (blue) and persulfidated (red) protein.

**Figure S8.**
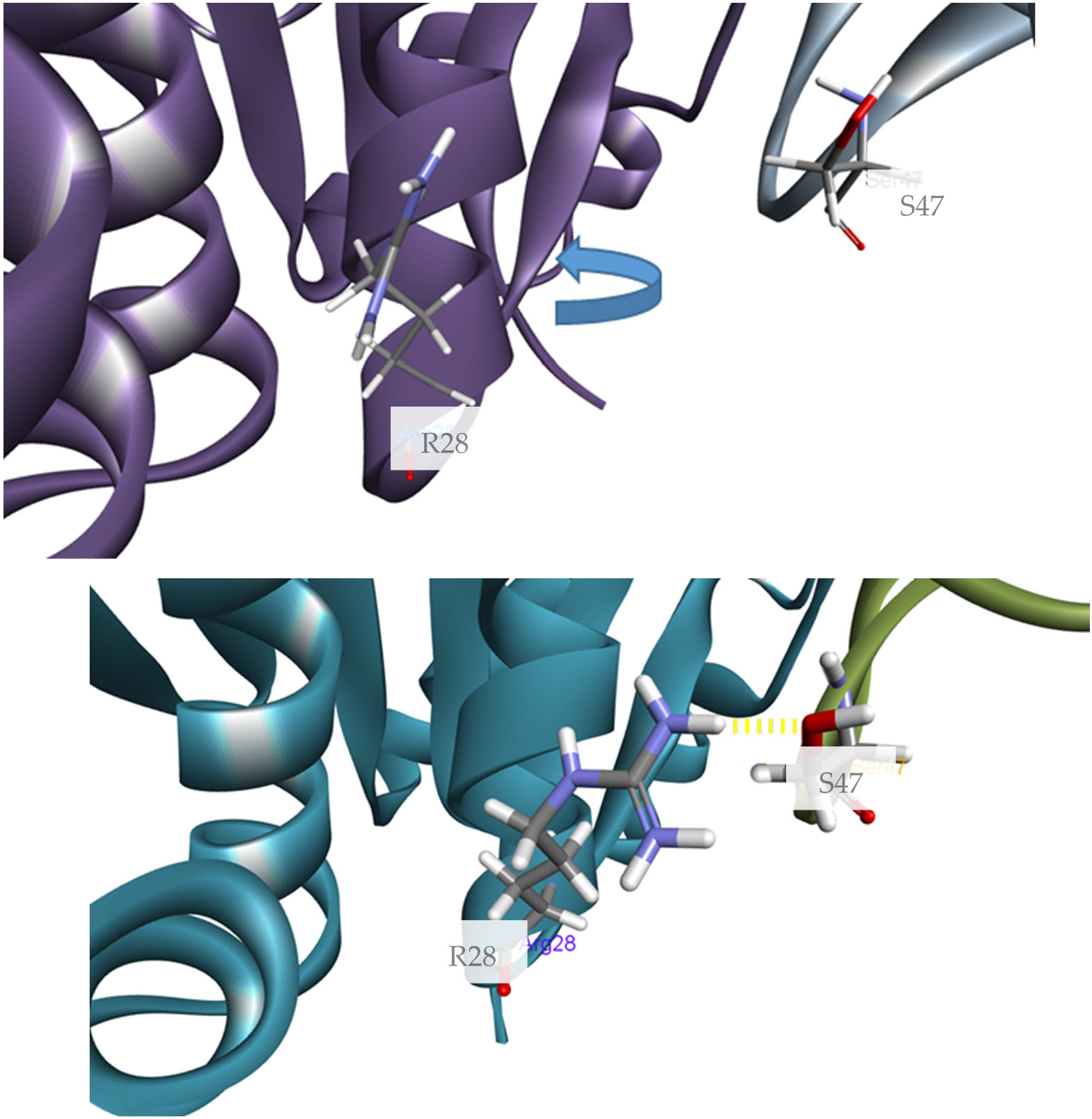
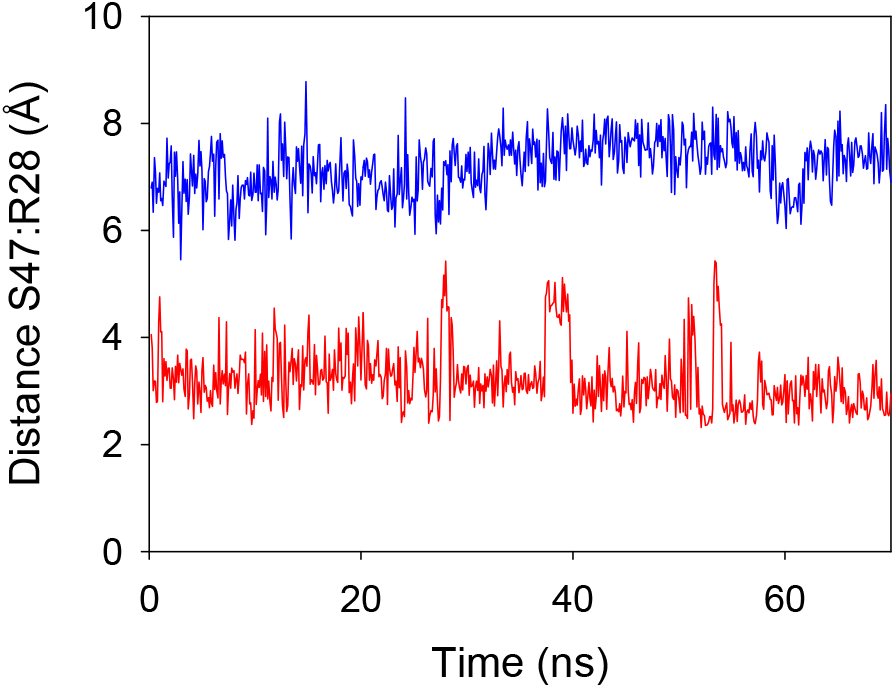
The interfacial interaction between S47:A and R28:B is strengthened after persulfidation. **A)** Representative snapshot obtained after 70 ns MD simulations of the interfacial interaction between S47 and R28, B) Time evolution of the distance between H-O group of S47 and the centroid of the NH_21/22_ from the guanidine group of R28 over the course of the simulation in the wt (blue) and persulfidated (red) protein.

**Figure S9.**
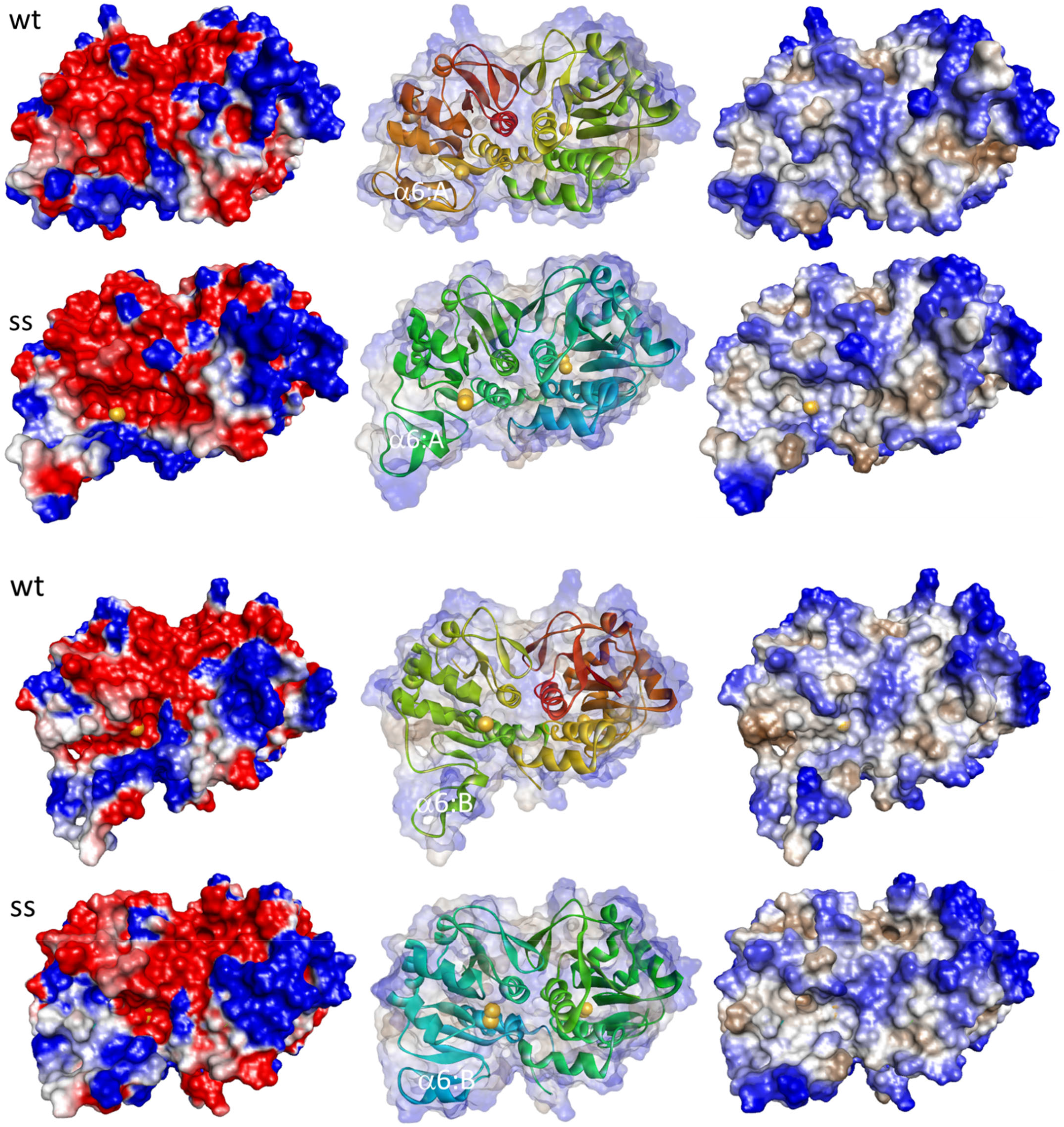
Persulfidation of DJ-1 induces changes at the protein surface. Electrostatic surface representation (left) and hydrophobicity surface (right) for both subunits of wt and SS-DJ1 after 70ns MD simulations. Electrostatic potential is colored to show negative charge in red and positive charge in blue. Hydrophobic and hydrophilic regions are representated in brown and blue, respectively. The localization of the protein subunits and of the sulfur atoms (yellow balls) are shown in the middle.

**Figure S10.**
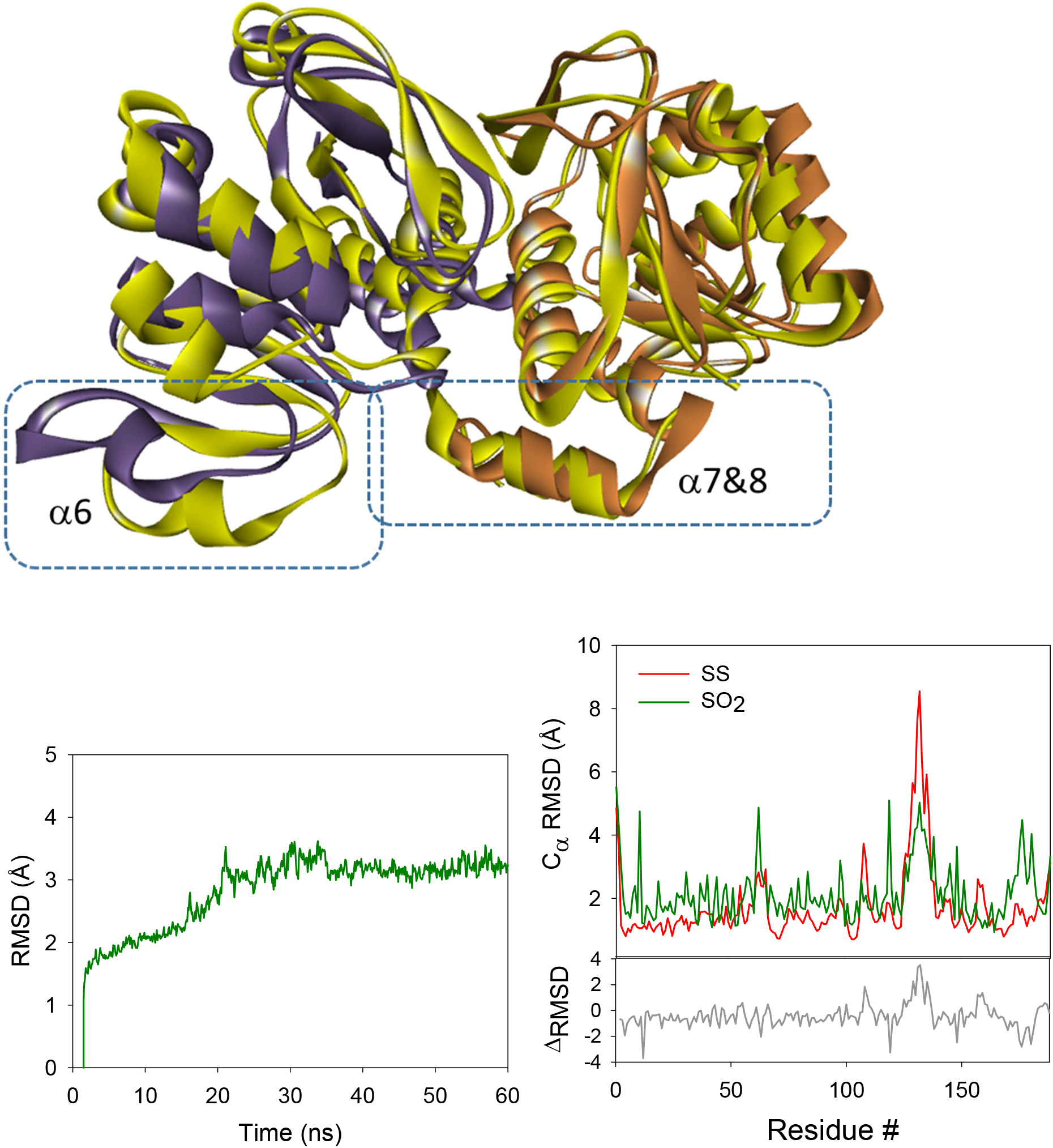
Structural comparison of sulfinylated and persulfidated DJ-1. A) Structural comparison of representative snapshots obtained after 60 ns MD simulations the persulfidated (yellow) and sulfinylated (purple and brown) DJ-1. The main region showing important differences in RMSD is highlighted. B) Root-mean-square deviation (RMSD) plot measured on the protein backbone backbone for the sulfinylated DJ-1 during the MD simulations with 100 ps interval for the sulfinylated DJ-1, and Cα-Root Mean Square Fluctuations (RMSD) plots for the persulfidated (red) and sulfinylated DJ-1 (green, average value of the individual values of the monomers). RMSD plots represent the average of the RMSD value of each monomer.

**Figure S11.**
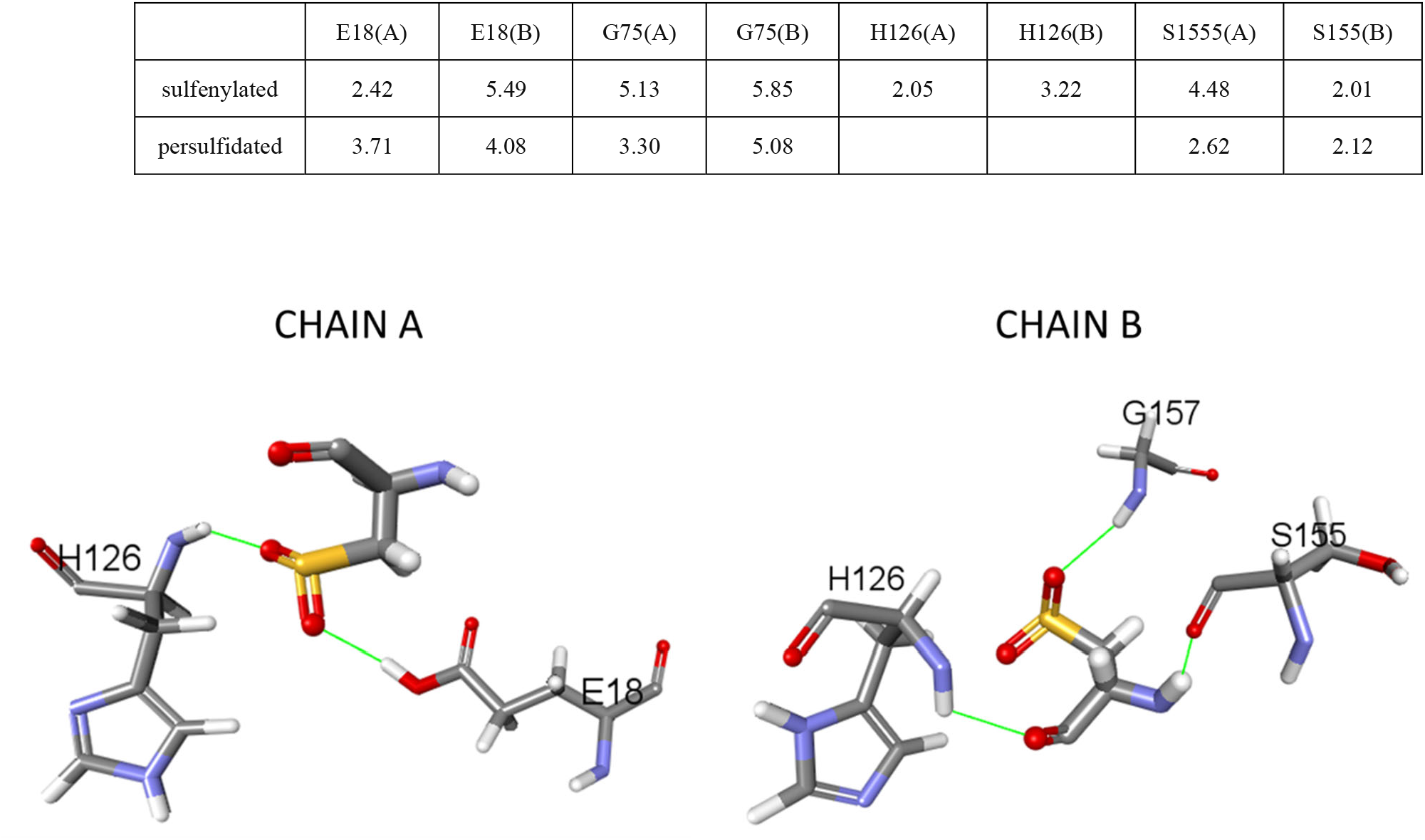
Main interactions between C106 and its neighbors in sulfinylated DJ-1. Average distances (Å) recorded over the last 45 ns of the dynamics between C106 and interacting residues. Distances are O-H(E18)…O=S(C106), N-H(G75)…O=C(C106), N-H(G75)…S(C106) (*), N-H(S155 or R156)…O=C(C106), N-H(H126)…O=C or O=S(C106). Views of the amino-acids interacting with C106.

**Figure S12.**
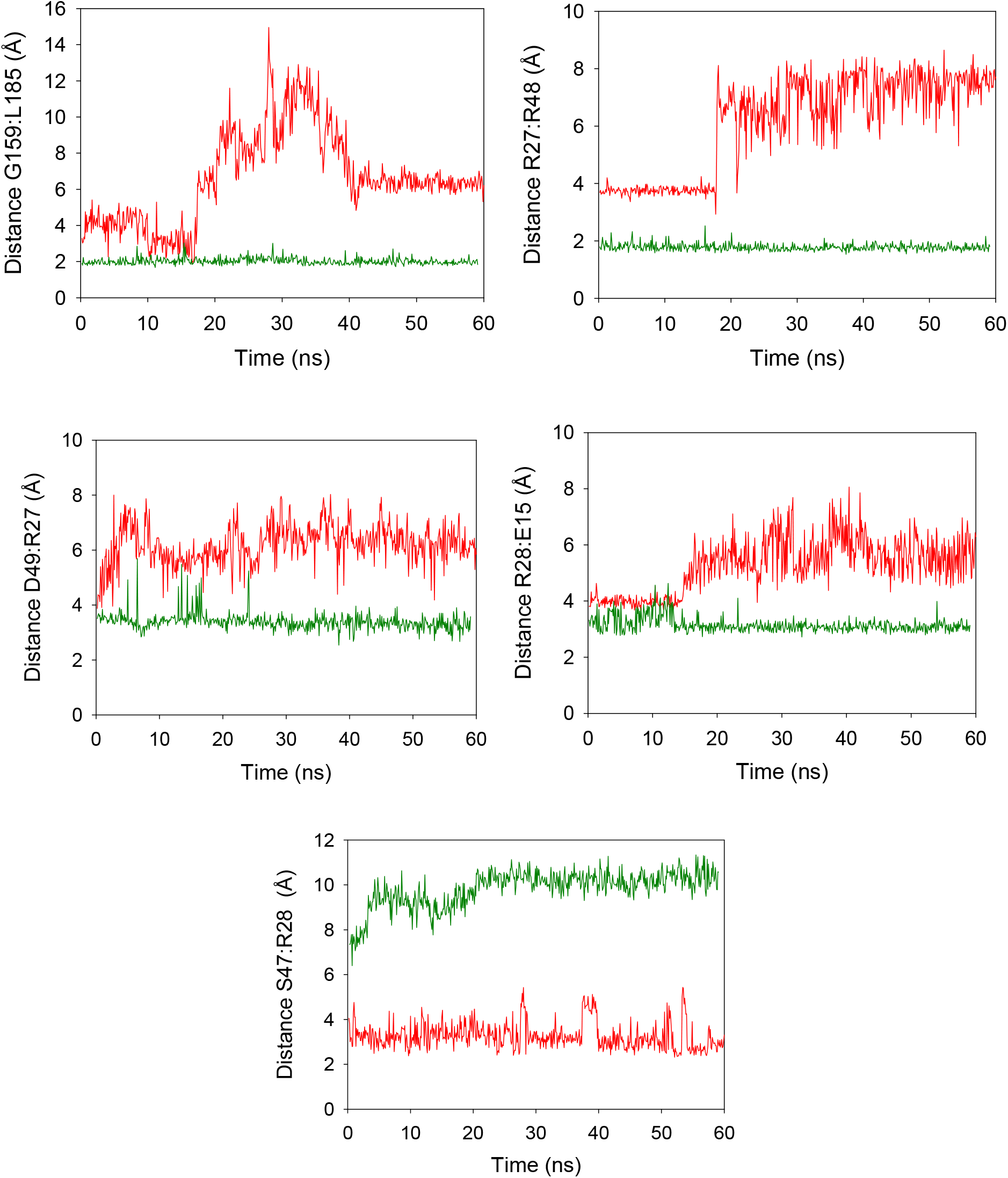
Differences in the representative interfacial contacts between the persulfidated and sulfinylated forms of DJ-1. Average distances (Å) recorded over the last 45 ns of MD simulations as described above for the persulfidated (red) and sulfinylated (green) forms of DJ-1.

**Figure S13.**
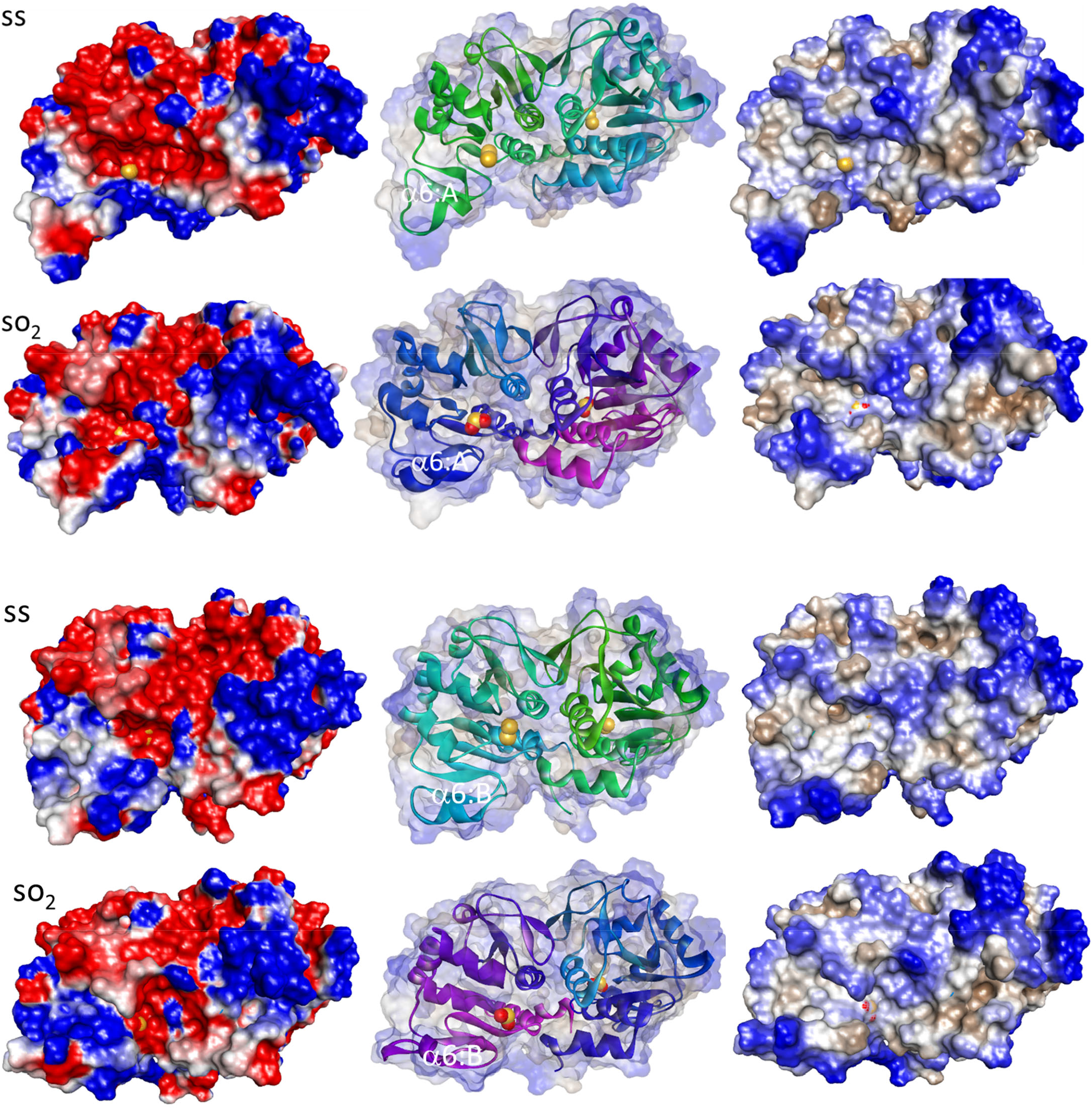
Persulfidation and sulfinylation induces different changes at the protein surface. Structural comparison of the surfaces from the wt, sulfinylated and persulfidated forms of DJ-1. Electrostatic surface representation (left) and hydrophobicity surface (right) for subunits A, B of wt and SS-DJ1 after MD simulations. Electrostatic potential is colored to show negative charge in red and positive charge in blue. Hydrophobic and hydrophilic regions are representated in brown and blue, respectively. The localization of the protein subunits and of the sulfur atoms (yellow balls) are shown in the middle.

